# Drug design and repurposing with a sequence-to-drug paradigm

**DOI:** 10.1101/2022.03.26.485909

**Authors:** Lifan Chen, Zisheng Fan, Jie Chang, Ruirui Yang, Hao Guo, Yinghui Zhang, Tianbiao Yang, Chenmao Zhou, Zhengyang Chen, Chen Zheng, Xinyue Hao, Keke Zhang, Rongrong Cui, Yiluan Ding, Naixia Zhang, Xiaomin Luo, Hualiang Jiang, Sulin Zhang, Mingyue Zheng

**Affiliations:** Drug Discovery and Design Center, State Key Laboratory of Drug Research, Shanghai Institute of Materia Medica, Chinese Academy of Sciences, 555 Zuchongzhi Road, Shanghai 201203, China; University of Chinese Academy of Sciences, No. 19A Yuquan Road, Beijing 100049, China; School of Chinese Materia Medica, Nanjing University of Chinese Medicine, 138 Xianlin Road, Jiangsu, Nanjing 210023, China; Shanghai Institute for Advanced Immunochemical Studies and School of Life Science and Technology, ShanghaiTech University, No. 393 Huaxia Middle Road, Shanghai 200031, China; The First Affiliated Hospital of USTC, Division of Life Sciences and Medicine, University of Science and Technology of China, 443 Huangshan Road, Hefei, Anhui 230027, China; Department of Analytical Chemistry, State Key Laboratory of Drug Research, Shanghai Institute of Materia Medica, Chinese Academy of Sciences, 555 Zuchongzhi Road, Shanghai 201203, China

## Abstract

Drug development based on target proteins has been a successful approach in recent decades. A conventional structure-based drug design pipeline is a complex, human-engineered pipeline with multiple independently optimized steps. Advances in end-to-end differentiable learning suggest the potential benefits of similarly reformulating drug design. Here, we proposed a new sequence-to-drug paradigm that discovers drug-like small-molecule modulators directly from protein sequences and validated this concept for the first time in three stages. First, we designed TransformerCPI2.0 as a core tool for the sequence-to-drug paradigm, which exhibited competitive performance with conventional structure-based drug design approaches. Second, we validated the binding knowledge that TransformerCPI2.0 has learned. Third, we applied a sequence-to-drug paradigm to discover new hits for E3 ubiquitin-protein ligases: speckle-type POZ protein (SPOP), ring finger protein 130 (RNF130) which does not have a 3D structure, and repurposed proton pump inhibitors (PPIs) for ADP-ribosylation factor 1 (ARF1). This first proof of concept shows that the sequence-to-drug paradigm is a promising direction for drug development.

## Introduction

Drug development based on proteins has been a successful approach in the past decades, for diseases with well-defined protein targets^1–3^. A typical protein structure-based drug design (SBDD) project starts from the protein sequence, builds a three-dimensional (3D) structure by structure biology or structure prediction, identifies binding pockets (orthosteric sites or allosteric sites), and finally discovers active modulators through virtual screening or de novo design^4,5^ (Fig. 1a). It involves a complex, human-engineered pipeline with multiple independently optimized steps, and each step has its own limitations^5^. First, a large number of proteins do not have high-resolution structures. Recently, the great success of AlphaFold^6^ and RoseTTAFold^7^ in protein structure prediction has given an overoptimistic view that the issue has been fully solved. However, not all the predicted structures are suitable for SBDD^8,9^, given that only 36% of all residues have very high confidence^10^. In particular, the precise structure prediction of active sites remains an unsolved challenge, since these local structures tend to break the ‘protein-folding rules’^9^. Second, prior knowledge about binding pockets is essential for SBDD, and it is a nontrivial issue to define such pockets for novel targets with multiple domains^11^. Moreover, predicting allosteric sites is still challenging^12^ due to the varied mechanisms of allosteric effects and high computational costs^13^. In addition, structural flexibility allows proteins to adapt to their individual molecular binders and undergo different internal motions^9,14,15^, making pockets more difficult to define. Third, virtual screening can produce false positives^16^ and may rapidly accumulate errors from the first two steps.

**Fig. 1.**
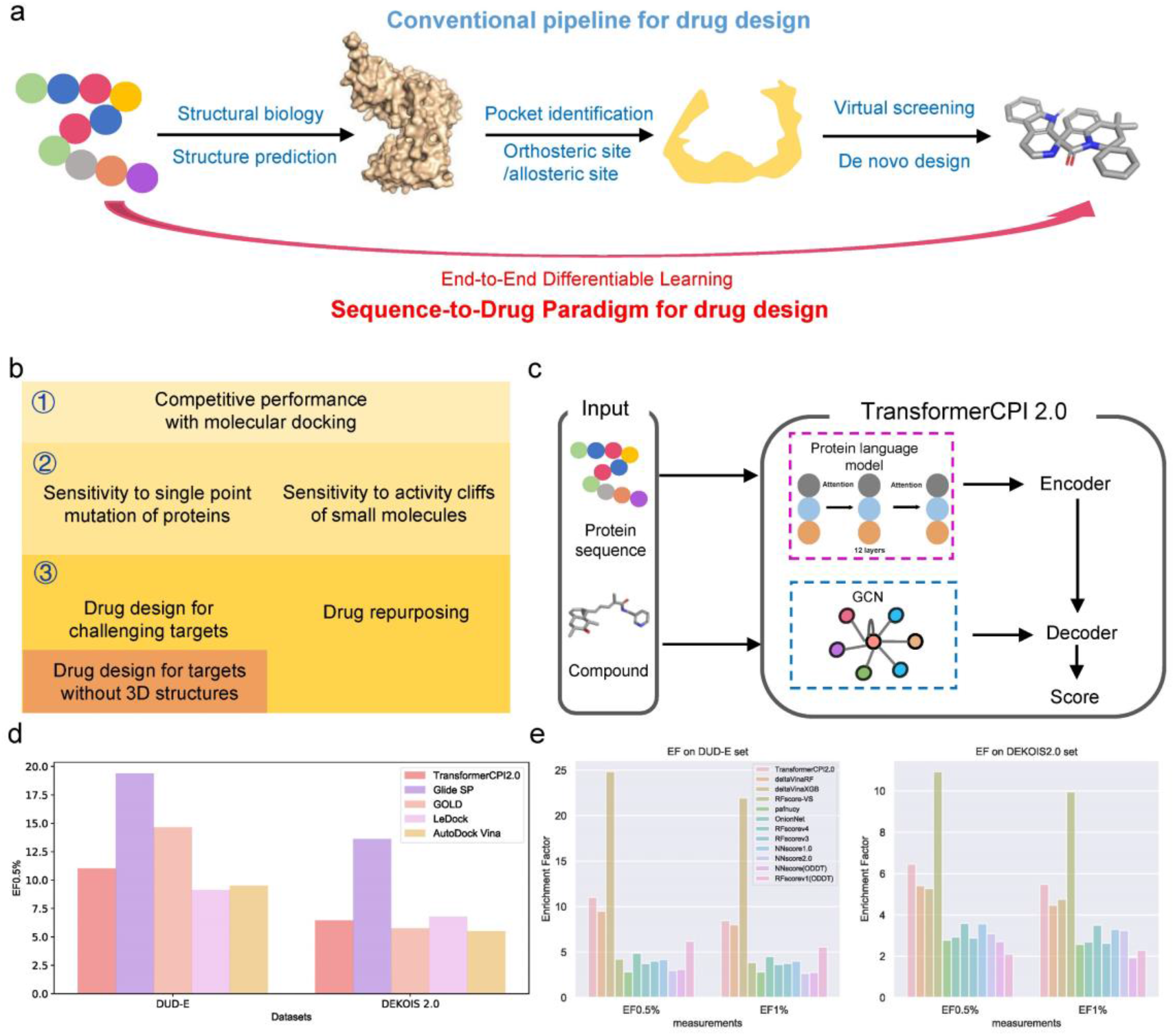
Competitive performance compared with 3D structure-based approaches. **a**, The conventional pipeline for target-based drug design and the sequence-to-drug paradigm. **b**, Three stages of the proof of concept of the sequence-to-drug paradigm, with each stage is labelled by different colours. **c**, The computational pipeline of TransformerCPI2.0. **d**, EF0.5% results of TransformerCPI2.0 and docking programs on DUD-E set and DEKOIS 2.0 set. **e**, EF0.5% and EF1% results of TransformerCPI2.0 and machine learning scoring functions on the DUD-E set and DEKOIS 2.0 set.

As an alternative to the conventional pipeline, we proposed a new sequence-to-drug paradigm that discovers modulators directly from protein sequences, jumping over the intermediate steps, by end-to-end differentiable learning (Fig. 1a). End-to-end differentiable deep learning has revolutionized computer vision and speech recognition^17^ by replacing all components of complex pipelines with differentiable primitives to enable joint optimization from input to output^18^. The success of AlphaFold^6^ also relies heavily on the idea of end-to-end differentiability. Therefore, this new paradigm is appealing because it performs the entire learning process in a self-consistent and data-efficient manner, avoiding the error accumulation of complex pipelines.

Many deep learning models have been proposed to use protein sequences as input^19–28^, but none have thoroughly verified the concept of the sequence-to-drug paradigm. In this work, we address the issue for the first time through three stages (Fig. 1b). First, we designed TransformerCPI2.0, which exhibited competitive performance against 3D structure-based approaches such as molecular docking, as a fundamental tool of the sequence-to-drug paradigm. Second, we tested whether TransformerCPI2.0 learns knowledge as expected, rather than exhibiting only data bias^20^. Limited effort has been devoted to investigating whether such a model learns in the manner we expected. For example, in the interaction between a receptor and a ligand, a tiny structural change in either the protein or the compound may lead to drastic changes in the interaction, but we do not know whether existing compound-protein interaction (CPI) models can capture such subtle changes and reflect them in the prediction results. Our previous analyses revealed that although some models achieved good performance on some datasets, they may not actually utilize protein-related information to make predictions^20^ and thus cannot generalize to mutants or new proteins. To judge whether a model truly has the ability to generalize, we need to examine whether it merely remembers the data distribution or learns the intermolecular recognition and regulation we are interested in. Therefore, we designed two computational analyses, i.e., drug resistance mutation analysis and molecular substitution effect analysis of the trifluoromethyl group, to validate the sensitivity of our model to single point mutations of proteins and activity cliffs of molecules. Third, we applied a sequence-to-drug paradigm to discover new hits for challenging targets, speckle-type POZ protein (SPOP) and ring finger protein 130 (RNF130) which does not have existing 3D structures. Additionally, we identified ADP-ribosylation factor 1 (ARF1) as a new target for proton pump inhibitors (PPIs). After the first proof of concept, the sequence-to-drug paradigm appears to be a promising direction for rational drug design.

## Results

### The sequence-to-drug paradigm achieves competitive performance against molecular docking

To build a model that can implement the sequence-to-drug paradigm, we developed TransformerCPI2.0 based on our previous work^20^, and its framework is shown in Fig. 1c. To train a general model for practical application, we constructed a chEMBL dataset containing 217,732 samples in the training set, 24,193 samples in the validation set and 10,199 in the test set. TransformerCPI^20^, CPI-GNN^19^, GraphDTA(GAT-GCN)^21^, and GCN^21^ were selected as baseline models, and all were retrained on the chEMBL dataset. We trained TransformerCPI2.0 and baseline models under the same criteria and compared their performance in terms of the area under the receiver operating characteristic curve (AUC) and the area under the precision recall curve (PRC) (Extended Data Fig. 1a∼c). TransformerCPI2.0 achieves the best performance among all the models. In addition, we tested TransformerCPI2.0 and the baseline models on the other three external datasets: GPCR set, Kinase set, and a large external set. TransformerCPI2.0 also showed the greatest generalization ability among all the models (Extended Data Fig. 1d∼i and Supplementary Table 1∼3). Furthermore, we designed a time-split test set named the chEMBL27 dataset to predict new data that were deposited online after the training set, and TransformerCPI2.0 still outperformed the baseline models (Extended Data Fig. 1j, k). Since our training set was generated from chEMBL23, this test suggested that our model can learn from past knowledge and generalize to future data.

We also compared TransformerCPI2.0 with conventional SBDD approaches for a better understanding of its capacity to enrich active molecules from large-compound collections. On two benchmark docking datasets, DUD-E^29^ and DEKOIS 2.0^30^, TransformerCPI2.0 achieved competitive performance against all the tested structure-based docking programs except Glide SP (Fig. 1d, Supplementary Table 4∼5). Furthermore, TransformerCPI2.0 yielded satisfactory performance compared with some well-known machine learning-based scoring functions (Fig. 1e, Supplementary Tables 6∼7). Overall, our method does not depend on the protein 3D structure yet achieves performance comparable to that of 3D structure-based molecular docking, and in subsequent studies, it will be used to solve virtual screening and target identification tasks.

### Sensitivity to drug resistance-associated single point mutations of protein

To further test whether TransformerCPI2.0 is sensitive to tiny local modifications of protein sequences, we proposed an analysis method named drug resistance mutation analysis, which mimics alanine scanning^31^. Briefly, we mutated each amino acid of the given protein sequence one by one and investigated whether the prediction score changed significantly. This analysis mimics the process of drug resistance in nature, where a single amino acid mutation may cause altered susceptibility to a drug.

We selected HIV-1 reverse transcriptase and its inhibitor doravirine as an example. Doravirine (formerly MK-1439) was approved by the FDA for the treatment of HIV-infected, treatment-naive individuals in combination with other antiretroviral drugs^32^. The cocrystal structure of HIV-1 reverse transcriptase and doravirine has been reported (PDB: 4NCG, Fig. 2a), and we conducted drug resistance mutation analysis and revealed the mutation effect of each amino acid position (Fig. 2c). It is encouraging that positions with a high relative activity change score (Δ*R*) are highly overlapped with the binding sites of doravirine (Fig. 2a, b), highlighting that TransformerCPI2.0 has implicitly learned information about binding site locations, since neither structure nor binding pocket information is included in the training phase. Among the mutations, P225 is predicted to be the most important site in HIV-1 reverse transcriptase (Fig. 2c) and actually plays an important role in the binding of doravirine (Fig. 2b). Additionally, P225, F227, L234 and P236 have been reported as drug resistance mutation sites^33–36^ and are correctly retrieved as important sites by TransformerCPI2.0. Furthermore, we analysed the specific mutation patterns predicted by TransformerCPI2.0 and found that some predictions matched the ground truth (Fig. 2d): P225H, F227C/L/R and P236L were correctly predicted by TransformerCPI2.0.

**Fig. 2.**
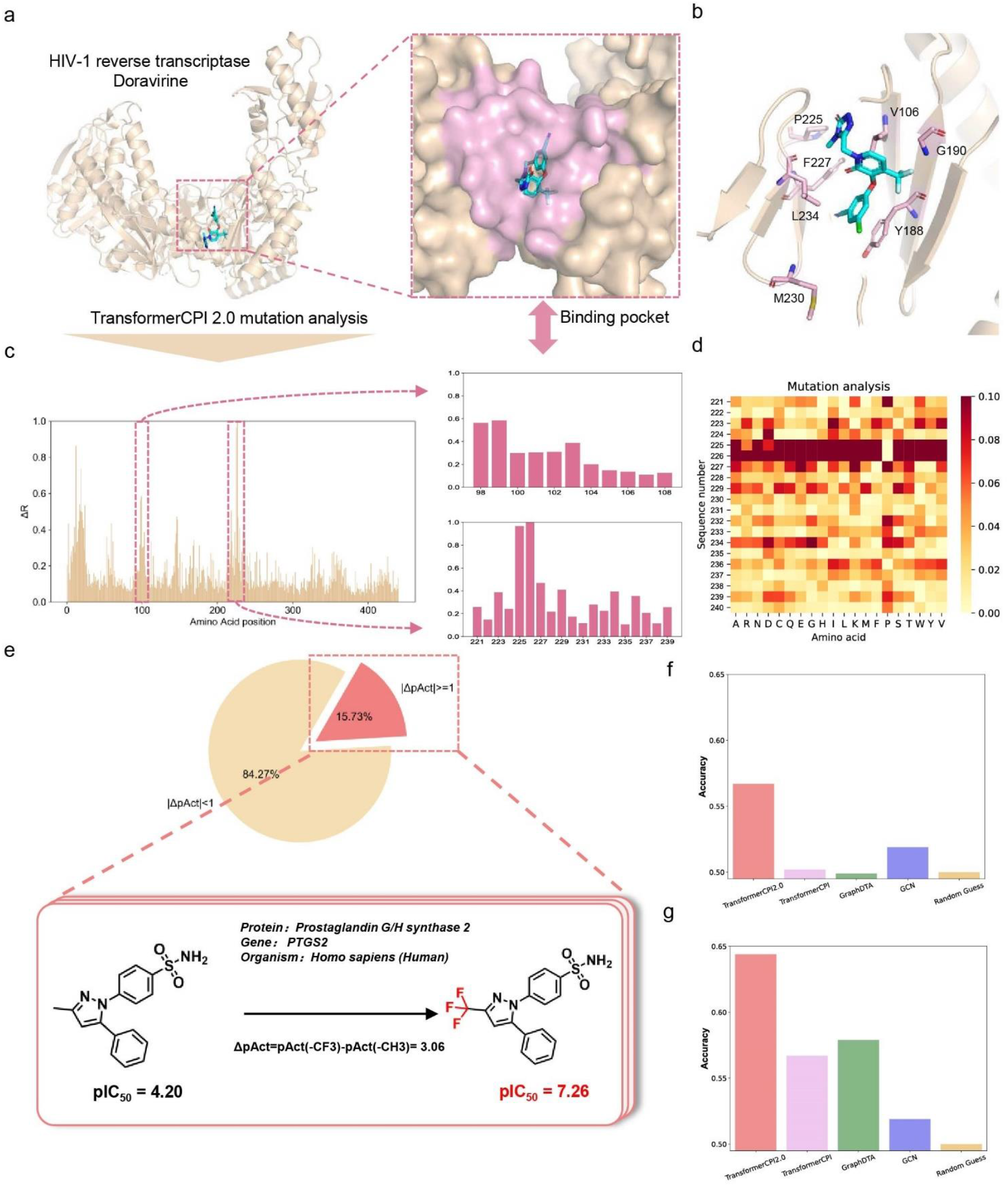
Drug resistance mutation analysis and substitution effect analysis of the trifluoromethyl group. **a**, The cocrystal structure of HIV-1 reverse transcriptase and doravirine (PDB: 4NCG). The binding pocket of doravirine is highlighted in pink. **b**, The binding mode of doravirine. The residues with drug-resistant mutations are coloured pink. **c**, Relative activity change score (Δ*R*) calculated by TransformerCPI2.0 at each amino acid position. The pink boxes mark the high Δ*R* regions, which are plotted in detail on the right. **d**, The heatmap plots the activity change score of positions 221 to 240, where each position is mutated to each of the 20 amino acids (including itself). The darker colour represents the higher activity change score caused by the mutation. **e**, Left, data distribution of the trifluoromethyl substitution dataset. Only 15.73% of substitution of −CH3 by −CF3 could increase or decrease the biological activity by at least an order of magnitude. We conducted substitution effect analysis on this part of data. Right, an example of −CH3 changed by −CF3 leads to a significant increase in biological activity. **f**, The accuracy of TransformerCPI2.0 and baseline models on the whole dataset. **g**, The accuracy of TransformerCPI2.0 and baseline models on the subset where the substitution of −CH3 by −CF3 could increase or decrease the biological activity by at least three orders of magnitude. **h**, An activity decrease example and an activity increase example are shown. The predictions of TransformerCPI2.0 are consistent with the ground truth for proteins and compounds not present in the training set.

Another possible concern is that the model might learn drug sensitivity-related residue information from the protein sequence alone rather than from the protein–ligand interactions that we expect it to learn^20^. We selected aspirin as a negative control (Extended Data Fig. 2a, b) and found that the pattern of Δ*R* was significantly different from that of doravirine. In conclusion, drug resistance analysis validated that TransformerCPI2.0 is sensitive to single point mutations of proteins and thus, similar to 3D structure-based docking methods, captures key residues for protein ligand binding.

### Sensitivity to activity cliffs of small molecules

To investigate whether TransformerCPI2.0 is sensitive to tiny modifications of ligands, we designed substitution effect analysis of the trifluoromethyl group as an example. Activity cliffs are generally understood as pairs or groups of similar compounds with large differences in potency^37,38^. Recently, Abula *et al*^39^ proposed a dataset including 18,217 pairs of compounds and corresponding bioactivity data, with the only difference being that −CH3 is replaced by −CF3. Only 15.73% of substitutions of −CF3 for −CH3 could increase or decrease the biological activity by at least an order of magnitude, and an example is shown (Fig. 2e).

We computed the activity change score Δ*s*_*c*_ and conducted substitution effect analysis on this part of the data. TransformerCPI2.0 reveals higher predictive ability than TransformerCPI, GraphDTA, GCN and random guessing (Fig. 2f and Supplementary Table 8). In addition, we investigated the performance of TransformerCPI2.0 and baseline models on a subset where substitution of −CH3 by −CF3 could increase or decrease the biological activity by at least three orders of magnitude. TransformerCPI2.0 still outperformed the other models, and the accuracy on this subset was higher than that on the whole dataset (Fig. 2g and Supplementary Table 9). The predictive capacity of this test is much more challenging than that of the whole dataset because the drastic biological activity change at this range involves turning an active compound into an inactive one or vice versa.

Furthermore, we chose illustrative examples to demonstrate the capacity of TransformerCPI2.0 (Extended Data Fig. 2c) to distinguish subtle structural differences that produce drastic activity changes, where none of the protein targets and compounds were in the training set. As shown, the activity change scores Δ*s*_*c*_ predicted by TransformerCPI2.0 are higher than 0.5. These four examples provide evidence that a single substitution in compounds will significantly change the prediction score of TransformerCPI2.0, which means that our model is sensitive enough to account for activity cliffs of small molecules.

### Drug design targeting E3 ubiquitin-protein ligases

SPOP, functioning as an adaptor of cullin3-RING ubiquitin ligase, mediates substrate protein recognition and ubiquitination^40,41^. A previous study validated SPOP as an attractive target for the treatment of clear-cell renal cell carcinoma (ccRCC), but it is a challenging target in terms of protein–protein interactions^42^. SPOP is overexpressed and misallocated in the cytoplasm of ccRCC cells, which induces proliferation and promotes kidney tumorigenesis^43^. Phosphatase and tensin homologue (PTEN) and dual specificity phosphatase 7 (DUSP7) are two substrates of SPOP^43^. PTEN acts as a negative regulator of the phosphoinositide 3-kinase/AKT pathway, and DUSP7 dephosphorylates extracellular signal-regulated kinase (ERK)^44^. Accumulation of cytoplasmic SPOP in ccRCC cells decreases cellular PTEN and DUSP7 by mediating the degradation of these two cytoplasmic proteins, leading to an increase in phosphorylated AKT and ERK and thus promoting the proliferation of ccRCC cells^43^.

Here, virtual screening with TransformerCPI2.0 was performed to discover new scaffold compounds that directly target SPOP (Fig. 3a, Supplementary Table 10) to test the feasibility of the sequence-to-drug paradigm on challenging targets. Four compounds were identified as initial hits by a fluorescence polarization (FP) assay (hit rate ∼5%), and 221C7 was the most active compound with an IC_50_ of 4.30 μM (Fig. 3b, c, Extended Data Fig. 3a, b). To further confirm that 221C7 disrupts SPOP-substrate interactions, an *in vitro* pull-down assay was performed. The results revealed that compound 221C7 dose-dependently reduced PTEN protein binding to the SPOP MATH domain (SPOP^MATH^) (Fig. 3d). Then, a nuclear magnetic resonance (NMR) experiment was conducted (Fig. 3e), and the result indicated direct binding between SPOP^MATH^ and 221C7. These results verified that 221C7 disrupts SPOP-substrate interactions by directly binding to SPOP^MATH^. Drug resistance mutation analysis was applied to interpret the prediction of TransformerCPI2.0, and important residues were mapped to the structure of full-length SPOP (PDB: 3HQI, Fig. 3f). Important residues mostly located on the MATH domain, and we then docked 221C7 to the predicted binding site (Fig. 3g).

**Fig. 3.**
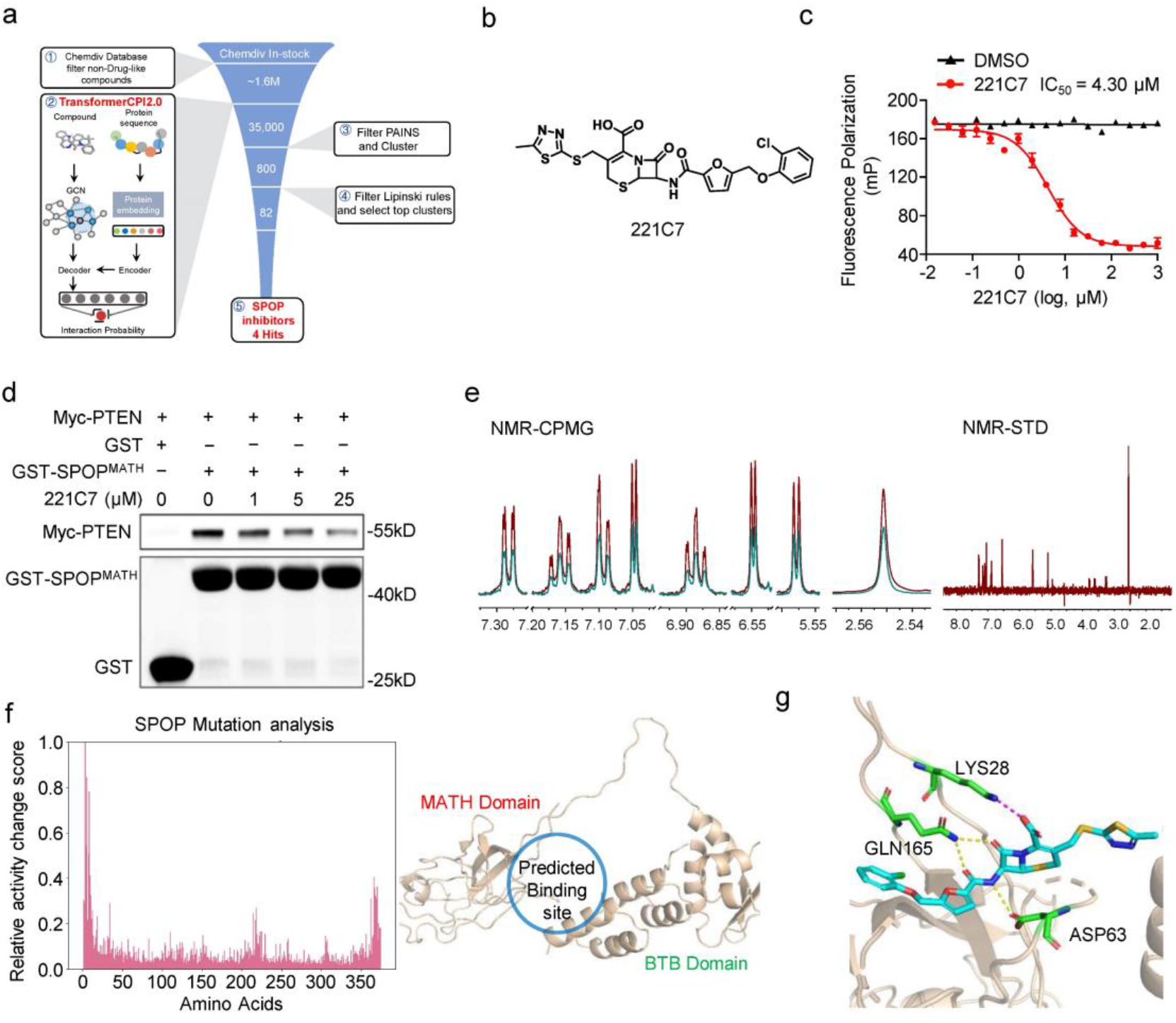
Discovering a novel scaffold hit of SPOP. **a**, Scheme of virtual screening protocol for small-molecule inhibitors of SPOP. **b**, Chemical structure of 221C7. **c**, 221C7 competitively inhibits puc_SBC1 peptide binding to SPOP^MATH^, as measured by the FP assay. **d**, 221C7 disrupts protein binding between SPOP^MATH^ and PTEN, as measured by *in vitro* pull-down assay. **e**, NMR measurement of direct binding between 221C7 and SPOP^MATH^. CPMG NMR spectra for 221C7 (red), 221C7 in the presence of 5 μM SPOP^MATH^ (green). The STD spectrum for 221C7 is recorded in the presence of 5 μM SPOP^MATH^. **f**, Drug resistance mutation analysis interprets the prediction of TransformerCPI2.0, and important residues are mapped back to the structure of full-length SPOP (PDB:3HQI). The predicted binding site is located on the MATH domain and near the BTB domain. **g**, The docking pose of 221C7 binding to SPOP (PDB: 3HQI).

The initial hit 221C7 was inactive in cellular experiments, which might be attributable to poor cell permeability caused by its large topological polar surface area (TPSA)^45^ of 214Å^2^. Therefore, we conducted hit expansion and obtained 26 structural analogues of 221C7, 19 of which were active in the FP assay (Extended Data Fig. 3c). Among them, 230D7 has a smaller TPSA (161Å^2^) and the smallest IC_50_ of the FP assay (Fig. 4a, b, Extended Data Fig. 3c), thus, it was selected for further validation. A protein thermal shift assay (PTS) revealed dose-dependent T_m_ shifts (Extended Data Fig. 4a), indicating that 230D7 could directly bind to SPOP^MATH^. Additionally, NMR experiments confirmed the direct binding between SPOP^MATH^ and 230D7 (Extended Data Fig. 4b). An *in vitro* pull-down assay was performed to verify that 230D7 dose-dependently reduces PTEN binding to SPOP^MATH^ (Extended Data Fig. 4c). After validating the molecular activity, we utilized 230D7 for the functional study at the cellular level.

**Fig. 4.**
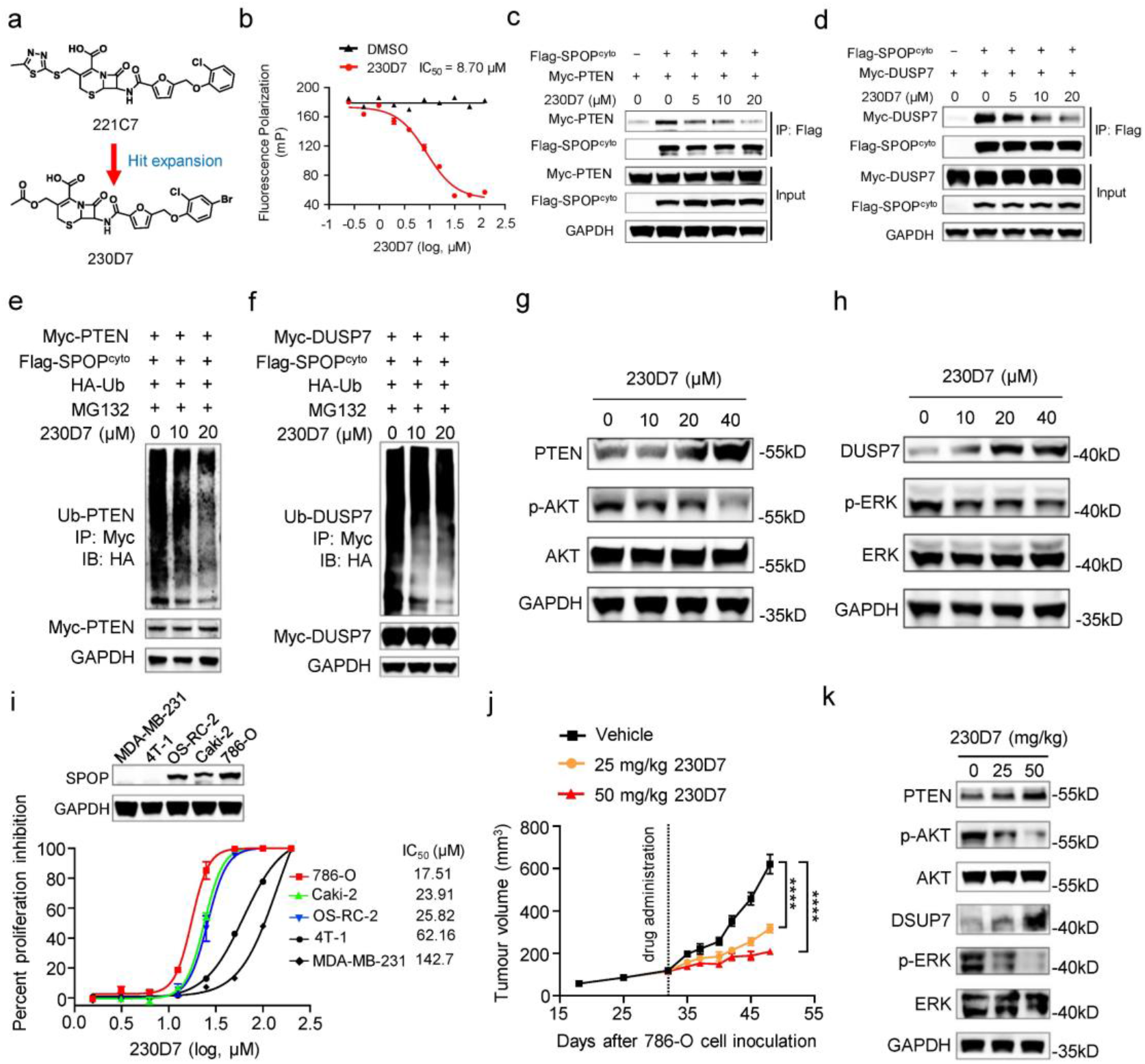
230D7 showed therapeutic potential for blocking oncogenic SPOP activity to treat ccRCC. **a**, Hit expansion of 221C7 to 230D7. **b**, 230D7 competitively inhibits puc_SBC1 peptide binding to SPOP^MATH^, as measured by the FP assay. **c∼d**, SPOP-PTEN and SPOP-DUSP7protein interactions are inhibited in the presence of 230D7 in 293T cells. **e∼f**, 230D7 inhibits the ubiquitination of PTEN and DUSP7 in 293T cells. **g∼h**, 230D7 upregulates PTEN and DUSP7 protein levels in 786-O cells. The downstream p-AKT and p-ERK abundances are observed to decrease. **i**, Cell proliferations of three ccRCC cell lines and two non-ccRCC cell lines in the presence of 230D7. The abundance of cytoplasm SPOP protein was measured. **j**, *In vivo* anti-ccRCC efficacy of 230D7 in 786-O xenograft models in NSG mice. Mice were administrated 230D7 at 25 or 50 mg/kg daily for 16 days by intraperitoneal dosing. Error bars represent mean ± SEM, with n = 7 for each group. P values were evaluated using 2-tailed unpaired t-test, ^****^P < 0.0001. **k**, Quantification of PTEN and DUSP7 accumulation and repression of p-AKT and p-ERK at day 16 of 786-O xenograft tumours treated with vehicle or 230D7.

We conducted a coimmunoprecipitation and *in vivo* ubiquitination experiment at the cellular level, and the results showed that 230D7 significantly disrupted the binding of PTEN and DUSP7 to SPOP in a dose-dependent manner (Fig. 4c, d), leading to decreases in PTEN and DUSP7 ubiquitination (Fig. 4e, f). Due to the inhibition of PTEN and DUSP7 ubiquitination under 230D7 treatment, accumulation of cellular PTEN and DUSP7 proteins was observed in 786-O cells treated with 230D7, causing decreases in phosphorylated AKT and ERK (Fig. 4g, h). Next, we tested the cell proliferation of three ccRCC cell lines (786-O, Caki-2, OS-CR-2) and two non-ccRCC cell lines (4T-1, MDA-MB-231) in the presence of 230D7 (Fig. 4i). 230D7 specifically inhibited the growth of ccRCC cell lines with an IC_50_ of approximately 20 μM compared with non-ccRCC cell lines. To determine if 230D7 is suitable for *in vivo* studies, we investigated the pharmacokinetics and acute toxicity profile of 230D7. 230D7 can be efficiently absorbed into the blood circulation following intraperitoneal injection and has low acute toxicity (Extended Data Fig. 4d∼h). A dose-dependent reduction in 786-O tumour growth rate could be observed in NSG mice treated with 230D7 (Fig. 4j), revealing significant anti-ccRCC therapeutic effect of 230D7 *in vivo*. No body weight loss in NSG mice was statistically noted during the entire pharmacodynamics study of 230D7 (Extended Data Fig. 4i). Finally, we checked the effect of 230D7 on oncogenic SPOP signalling in ccRCC xenograft tumours. As expected, PTEN and DUSP7 are elevated in 230D7 treated groups, and p-AKT and p-ERK levels are decreased (Fig. 4k). Moreover, we confirmed the high selectivity of 230D7 which does not target kinases (Extended Data Fig. 5). In conclusion, we successfully discovered novel small-molecule inhibitors that target SPOP through a sequence-to-drug paradigm, among which 230D7 showed therapeutic potential for blocking SPOP activity to treat ccRCC.

After discovering novel inhibitors for SPOP, we applied this paradigm to discover hits for a more challenging target RNF130 whose crystal structure is unknown. RNF130 is an E3 ubiquitin-protein ligase without structural information, and no chemical binders have been reported. Our recent study revealed that RNF130 plays an important role in autoimmune inflammation, suggesting its inhibition could be of potential therapeutic value (unpublished data, under review). We utilized TransformerCPI2.0 to screen compounds that directly bind to RNF130 (Extended Fig. 6a, Supplementary Table 11) and discovered iRNF130-63 to be a binder of RNF130 (Extended Fig. 6b∼d). The direct binding between iRNF130-63 and RNF130 protein was confirmed through surface plasmon resonance (SPR), and this binding exhibited a fast-on, fast-off kinetic pattern with a K_D_ of 9.36 μM (Extended Fig. 6c). We also performed a cellular thermal shift assay (CETSA), and the results supported that iRNF130-63 directly binds to and thermally stabilizes the RNF130 proteins (Extended Fig. 6d). The success of discovering novel hits for SPOP and RNF130 demonstrated that the sequence-to-drug paradigm is practicable for virtual screening with encouraging prospects.

### Repositioning proton pump inhibitors as anticancer drugs by targeting ARF1

Benefiting from the end-to-end nature, the sequence-to-drug workflow can be inversely used to enable drug target identification or drug repurposing. This means that we can perform proteome-wide target screening, as only protein sequence information is required except for a given drug molecule as the model input. Here, we select proton pump inhibitors (PPIs) as a drug repurposing case study. To date, preclinical data and clinical data support the use of PPIs in cancer treatment^46^, but few new targets have been found. TransformerCPI2.0 was applied to score 2,204 human proteins from the DrugBank database^47^ against four classical PPIs (rabeprazole, lansoprazole, omeprazole and pantoprazole, Fig. 5a∼b, Supplementary Table 12), and the results were sorted by predicted interaction probability. After analysing the top 20 proteins, ARF1 attracted our attention due to its oncogenic effect on cancer stem cells (CSCs) by the lipolysis pathway^48,49^. ARF1 is a small G protein and belongs to the RAS superfamily, which cycles between an active GTP-bound and an inactive GDP-bound conformation^50^. Recent studies have shown that the lipid metabolism regulated by ARF1 selectively maintains cancer stem cells (CSCs), and ARF1 inhibition or knockdown in CSCs leads to the accumulation of lipid droplets, further leading to metabolic stress, which can not only selectively kill CSCs but also stimulate an anticancer immune response and achieve lasting therapeutic effects^48,49^. Inhibition of ARF1 activity is a promising direction for cancer immunotherapy, therefore, we selected ARF1 for investigation.

**Fig. 5.**
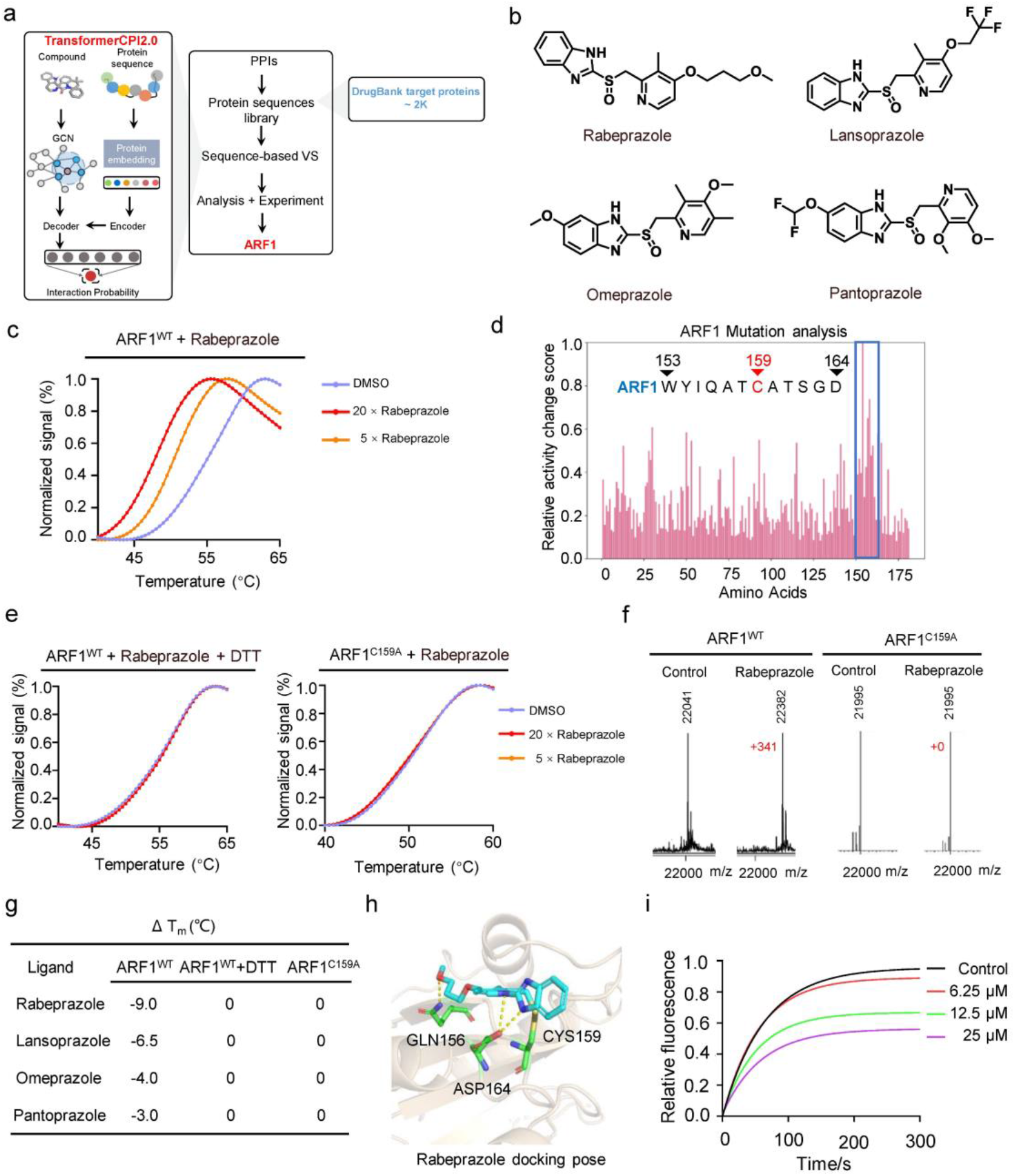
Identifying ARF1 as the new target of PPIs. **a**, Scheme of target identification protocol for PPIs. **b**, Chemical structures of four PPIs, rabeprazole, lansoprazole, omeprazole and pantoprazole. **c**, Effect of rabeprazole on the thermal stability of ARF1^WT^ in the PTS assay. **d**, Drug resistance mutation analysis interprets the prediction of omeprazole, and there is a cysteine (C159, the only cysteine residue of ARF1) residue among the important residues. **e**, Effects of rabeprazole on the thermal stability of ARF1^WT^ (containing DTT) and ARF1^C159A^ in the PTS assay. **f**, The protein molecular weights of ARF1^WT^ and ARF1^C159A^ in the presence or absence of rabeprazole were determined by mass spectrometer. **g**, Summary of PTS assay results. **h**, Potential docking pose of rabeprazole and ARF1 (PDB: 1HUR). **i**, GDP/MANT-GTP nucleotide exchange of ARF1 treated with rabeprazole.

PTS assay revealed dose-dependent T_m_ shifts (Fig. 5c, Extended Data Fig. 7b∼d), indicating that PPIs could directly bind to wild-type ARF1 (ARF1^WT^) and destabilize the protein. Drug resistance mutation analysis was then applied to interpret the prediction of TransformerCPI2.0. The results indicated that amino acids 150∼165, a range that contains a cysteine (C159), contributed greatly to compound protein binding (Fig. 5d, Extended Data Fig. 7a). Given that PPIs covalently bind to the cysteine of H^+^/K^+^-ATPase^51^, we conducted different assays to determine whether PPIs covalently bind to ARF1. Therefore, another two PTS assays were conducted: 1) PPIs with ARF1^WT^ and the reducing agent dithiothreitol (DTT), which can break disulfide bonds; and 2) PPIs with ARF1^C159A^ where C159 was mutated to alanine. No T_m_ shifts were observed in neither two assays (Fig. 5e, Extended Data Fig. 7b∼d). Mass spectrometry (MS) further validated that PPIs can covalently bind to ARF1^WT^ but cannot bind to ARF1^C159A^ (Fig. 5f, Extended Data Fig. 7e∼f), and two-dimensional mass spectrometry determined the covalent binding site is C159 (Extended Data Fig. 8a∼d). Among the four PPIs, rabeprazole had the greatest effect on the thermal stability of ARF1 (Fig. 5g), thus we selected rabeprazole for further functional study. According to the covalent binding site, we provided a possible docking pose of rabeprazole (Fig. 5h). Activation of ARF1 requires the release of GDP followed by the binding of GTP, a process catalysed by guanine nucleotide exchange factor (GEF)^50^. Therefore, we performed GDP/MANT-GTP nucleotide exchange catalysed by ARNO (a type of GEF) and found that rabeprazole suppressed the nucleotide exchange process in a concentration-dependent manner (Fig. 5i), verifying its inhibition of ARF1 activity.

According to previous works^48,49^, we first detected the inhibitory effect of rabeprazole on the activity level of ARF1 in CT26 cells (colon carcinoma cells) using a G-LISA assay. The results showed that rabeprazole effectively inhibited ARF1 activity in CT26 cells in a concentration-dependent manner (Fig. 6a). In addition, significant accumulation of lipid droplets was observed in CT26 cells treated with rabeprazole (Fig. 6b). To evaluate the antitumour effect of rabeprazole *in vivo*, we established colon cancer transplanted tumour models by injecting CT26 cells into BALB/c mice. Rabeprazole treatment significantly suppressed the tumour growth in mice as measured by tumour volume (Fig. 6c). In order to verify that rabeprazole induce an antitumour immune response, we analysed the immune cell subsets of colon cancer transplanted tumours by fluorescence-activated cell sorting (FACS) and found a significant increase in CD3^+^ CD8^+^ T cells, as well as a significant decrease in CD3^+^ CD8^+^ PD1^+^ T cells, CD3^+^ CD8^+^ TIM3^+^ T cells and CD3^+^ CD8^+^ PD1^+^TIM3^+^ T cells (Fig. 6d). Additionally, upregulation of CD8 and downregulation of PD1 were detected by immunohistochemical staining (Fig. 6e), confirming that an antitumour immune response was stimulated by rabeprazole. All these data suggest that rabeprazole inhibits the growth of colon cancer by inducing an antitumour immune response. In summary, the success of repurposing PPIs to ARF1 demonstrated that the inverse application of the sequence-to-drug paradigm for drug repositioning is also practicable with encouraging prospects.

**Fig. 6.**
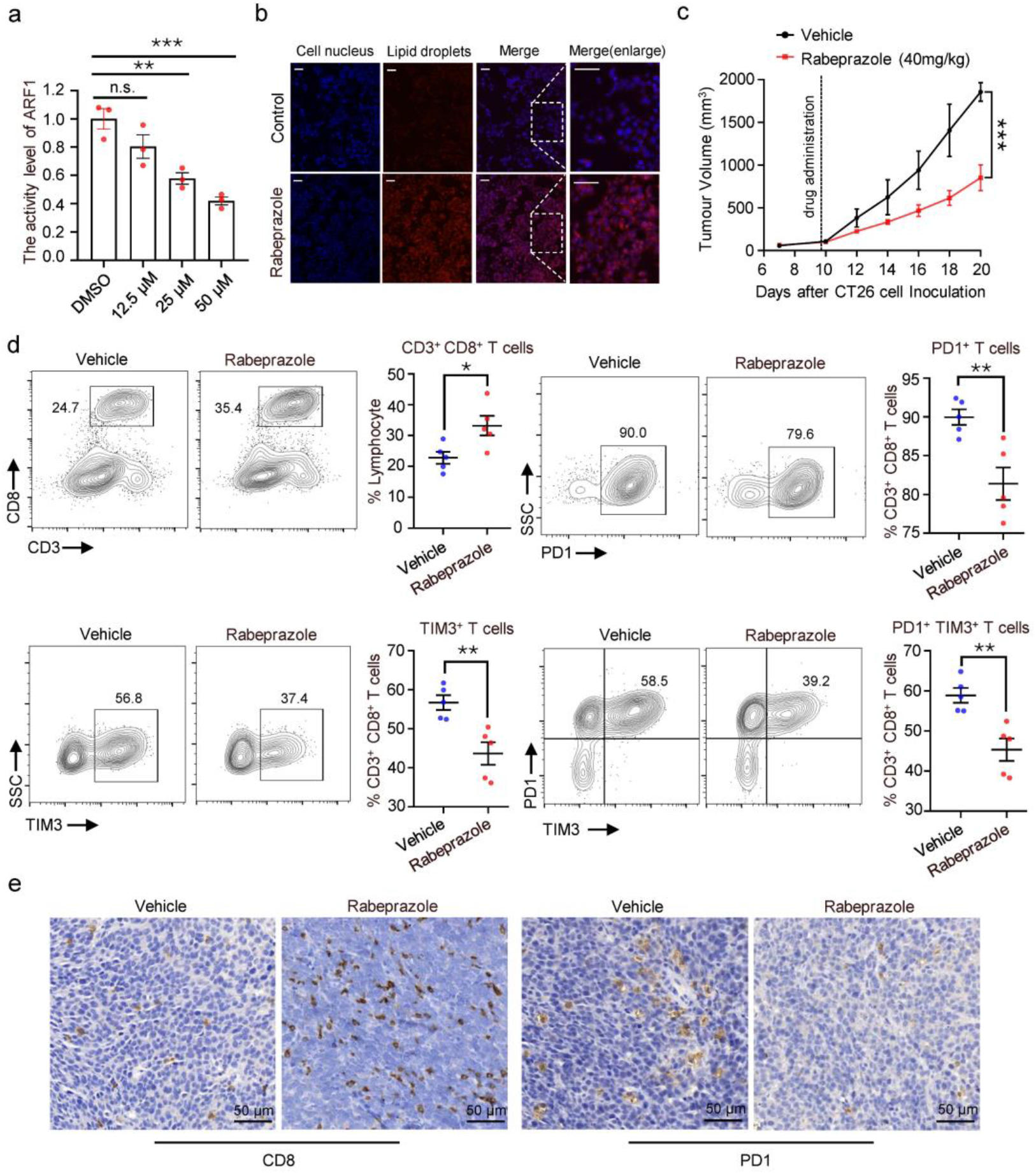
Rabeprazole induces antitumour immune response through lipid metabolism. **a**, The activity level of ARF1 in CT26 cells treated with rabeprazole for 48h was measured by using G-LISA assay. **b**, Fluorescent images of CT26 cells stained with DAPI (for nucleus) or Nile red (for lipid droplets) after treatment with rabeprazole (20 μM) or DMSO. Scale bars: 100 μm. **c**, *In vivo* efficacy of rabeprazole in CT26 transplanted tumour model in BALB/c mice. Mice were administrated rabeprazole 40 mg/kg daily for 10 days by intraperitoneal dosing. **d**, Impact of rabeprazole delivery on immune cell subsets in CT26 transplanted tumour model, assessed by flow cytometry analysis (a, c-d, Error bars represent mean ± SEM, with n = 5 for each group. P values were evaluated using 2-tailed unpaired t-test. ^*^P < 0.05, ^**^P < 0.01, ^***^P < 0.001). **e,** Immunohistochemical staining for cell surface markers (CD8, PD1) of tumour tissues in the indicated groups.

## Discussion

Conventional structure-based drug design pipeline is a complex, human-engineered pipeline with multiple independently optimized steps. Structural functional insights and profound experience are needed to build 3D structures through structural biology or structure prediction, identify the potential binding pockets, either orthosteric sites or allosteric sites, and screen active molecules by different molecular docking protocols. However, such multistep operation is error prone; for example, inaccurate protein structures, the multiplicity and dynamics of binding pockets, incorrect pocket definition, inappropriate selection of the scoring function, etc., and the errors or intrinsic accuracy limitations of each step accumulate rapidly and significantly lower the success rate. When little information about target proteins is known, this issue becomes more serious and has formed a long-lasting obstacle to rational drug design.

To address this problem, we proposed a new sequence-to-drug paradigm that discovers modulators directly from protein sequences, jumping over intermediate steps. This proof-of-concept study analyses the sequence-to-drug paradigm in three aspects: its competitive performance against 3D structure-based molecular docking, its learning of binding knowledge, and drug design for challenging targets and drug repurposing. Apart from methodology, the sequence-to-drug paradigm successfully discovered novel inhibitors for SPOP, among which 230D7 showed potential to treat ccRCC, and discovered the first binder of RNF130. Given that SPOP is an adaptor of E3 ligase and RNF130 is an E3 ligase, the hits we found have the potential to serve as novel warheads for proteolysis-targeting chimaeras (PROTACs). PROTACs have been successfully developed for harnessing the ubiquitin–proteasome system to degrade a protein of interest, receiving tremendous attention as a new and exciting class of therapeutic agents that promise to significantly impact drug discovery^52^. Due to the limited structural knowledge of E3 ligases and their substrate recognition and regulation of ubiquitination, the hits we discovered may promote the development of PROTACs. Furthermore, through an inverse application of the sequence-to-drug paradigm, the FDA-approved drug rabeprazole showed promise for expanding its indications to colon cancer treatment by regulating lipid metabolism and inducing an antitumour immune response. All these results demonstrate that the sequence-to-drug paradigm is a promising direction for drug design.

Recently, due to the rapid growth of virtual libraries, such as DrugSpaceX^53^ and synthesis (REAL) combinatorial libraries^54^ which cover spaces of 100 million to multiple billions of chemicals, there is a high demand for developing computationally efficient virtual screening approaches. The sequence-to-drug paradigm can be combined with these large virtual libraries to rapidly discover new active scaffolds from the unexplored chemical space. On the one hand, the explosion in available genomic sequencing techniques and annotation techniques has revolutionized bioinformatics^55^, and the sequence-to-drug paradigm will likewise evolve rapidly by incorporating more abundant multiple sequence alignment or functional annotation information. On the other hand, we may perform more comprehensive proteome-wide virtual screening, accelerating the discovery of new drug hits for novel but challenging biological targets. Overall, our findings provide a proof of concept for the sequence-to-drug paradigm, which we believe will become an essential component of future rational drug design pipelines.

## Method

### TransformerCPI2.0 model and training details

Compared with TransformerCPI, TransformerCPI2.0 has been updated in the following four aspects: (1) removing 3-gram protein word embedding calculated by the Word2Vec algorithm, (2) computing protein sequence representation by a pretrained protein language model named TAPE-BERT, (3) replacing 1D convolutional neural networks and gated linear units with a self-attention-based transformer encoder, and (4) introducing a new atom vector into the atom sequence that carries the interaction information at the molecular level.

### Pretraining the protein language model

Word2vec is an unsupervised technique to learn high-quality distributed vector representations that describe sophisticated syntactic and semantic word relationships and maps discrete words to low-dimensional real-valued vectors. However, the final embedding table of Word2vec is stationary, regardless of the upstream and downstream context information of the given word, which may lead to errors regarding the true meaning of the word in its local context. Since BERT^56^ achieved great success in natural language processing (NLP), many efforts have been devoted to protein sequence representation learning. Many pretraining models based on long short-term memory (LSTM) or transformer architectures have been proposed, such as UniRep^57^ and TAPE^58^. To maintain the model consistency and gain parallel computing efficiency, we chose the transformer model in TAPE (TAPE-BERT) to calculate protein sequence embedding. The TAPE-BERT model contains 12 self-attention encoder layers, 12 attention heads for each layer, 768 dimensions for the hidden state, and 3072 dimensions for feedforward layers. We first utilized the TAPE Tokenizer from TAPE-BERT to encode the protein amino acid sequence into real values, where numbers 1∼23 represent 23 common amino acids, 0 represents the token ‘<PAD>’, 24 represents the token ‘<CLS>’, and 25 represents the token ‘ <SEP>’. Then, we input this encoded real value sequence into the pretrained TAPE-BERT model and finally obtained protein embeddings with 768 dimensions. In TransformerCPI2.0, protein embedding from the TAPE-BERT model serves as the input to the encoder of TransformerCPI2.0, replacing the embedding calculated from the Word2Vec model.

### Encoder of TransformerCPI2.0

Since protein embedding was calculated by the TAPE-BERT model, we replaced 1D convolutional neural networks and gated linear units with the original self-attention-based transformer encoder. Given that the position information of the amino acid sequence has been taken into consideration when computing protein embeddings, position embedding was removed from the transformer encoder. The encoder consists of 3 encoder layers, 8 attention heads for each layer, 768 dimensions for the hidden state, and 3072 dimensions for feedforward layers. To maintain the maximal performance of TAPE-BERT, the hidden state dimension of the encoder was exactly the same as that of TAPE-BERT, ensuring no information loss in this process. After the hyperparameter search, 12 attention heads showed little performance improvement compared to 8 attention heads but much higher training and inference time; therefore, only 8 attention heads were used in TransformerCPI2.0.

### Atom embedding calculation

Each of the atom features was initially represented as a vector of size 34 using the RDKit python package, and the list of atom features can be found in our previous work. In TransformerCPI2.0, we additionally introduced a new virtual atom that carries the information at the molecular level and does not exist in the given compound. This virtual atom was initialized as the average of atom features across the whole compound and was linked to all atoms. All the atom vectors together with the virtual atom vector were put into graph convolutional networks (GCNs)^59^ to learn the representation by integrating their own neighbourhood information. Notably, only one GCN layer was used and recommended in this process, and more than two GCN layers harmed the performance of TransformerCPI2.0 to a great deal. Too many GCN layers may over smooth the atom features, causing different atom features to tend to be similar to each other. TransformerCPI2.0 then fails to learn compound-protein interaction features when the atom embedding carries excessively similar features.

### Decoder of TransformerCPI2.0

Protein embedding and atom embedding serve as the target sequence and memory sequence of the transformer decoder, respectively. Consistent with the encoder, the decoder consists of 3 decoder layers, 8 attention heads for each layer, 768 dimensions for the hidden state, and 3072 dimensions for feedforward layers. In addition, the original transformer was designed to solve seq2seq tasks and utilize a causal mask operation to cover the downstream context of each word in the decoder. We removed the mask operation of the decoder to ensure that our model accesses the whole target sequence. Since we introduced a new virtual atom, as described above, we used the last layer representation of this virtual atom rather than the weighted sum of the last layer atom representation to predict the compound protein interaction probability. The last layer presentation of virtual atoms was fed into fully connected layers and finally returned the compound protein interaction probability.

### Training details

The TransformerCPI2.0 model was trained by the RAdam^60^ optimizer with a learning rate of 1e-5 and a weight decay of 1e-3. The batch size of 1 was selected to ensure that the longest protein sequence fit into the GPU memory. We employed the gradient accumulation technique to expand the actual batch size to 64. Training was performed on one NVIDIA Tesla V100 (16G) GPU. The TransformerCPI2.0 model was trained for ∼50 epochs or ∼1.5 weeks of wall clock time.

### ChEMBL dataset construction

Inheriting our previous work, we followed two rules to construct a dataset: (i) CPI data was collected from an experimentally validated database and (ii) each ligand should exist in both positive and negative classes. To build a universal deep learning model for all types of proteins, we selected the chEMBL_23^61^ database to construct a universal dataset to train TransformerCPI2.0.

### Data cleaning

The ChEMBL_23 database was released on 1 May 2017 and contains 2,101,843 compound records, 1,735,442 compounds, 14,675,320 activities, 1,302,147 assays, 11,538 targets and 67,722 source documents. We downloaded the whole database and cleaned the data using the following procedure:

1. The target type was set to ‘SINGLE PROTEIN’, and the molecule type was set to ‘Small molecule’;
2. Data with a confidence score of 9 and assay type of ‘B’ were reserved;
3. Activity data with activity metrics of IC_50_, EC_50_, K_i_ in units of nM were selected.

### Dataset process

After cleaning the data, we transformed IC_50_, EC_50_, and K_i_ to pIC_50_, pEC_50_, and pK_i_ and then split the dataset into a positive set and a negative set at the threshold of 6.5. Samples with different labels were removed from the dataset. In the early stage of drug discovery, only hit compounds whose IC_50_, K_i_ or EC_50_ are at the μM or even nM level will be further optimized. Additionally, public data are prone to a certain experimental error, i.e., on average 0.5 log units for IC50 data^62,63^. The threshold of 6 considers those CPI pairs whose IC50, Ki or EC50 are smaller than 1 μM as positive samples, which causes the models to select CPI pairs with high activity. To decrease the risk of experimental error in public data, a stricter threshold of 6.5 was used in a previous work^64^. Therefore, we selected the threshold of 6.5 to define positive data and negative data. Data whose atom number was more than 60 or whose protein sequence length exceeded 4000 were filtered out, guaranteeing that all the data fit into GPU memory. Finally, we constructed a ChEMBL dataset including 3,348 proteins, 69,616 compounds, 117,513 positive CPIs, 134,611 negative CPIs and 252,124 samples in total.

### Dataset split and label reversal experiment

We selected ligands that exist in both the positive and negative classes to make the compound distribution in positive samples and negative samples exactly the same, trying to eliminate the potential ligand bias as much as possible. Consequently, we used a label reversal experiment to split the ChEMBL dataset. For the ChEMBL dataset, we randomly selected 2,941 ligands and pooled all the negative CPI samples containing these ligands into the test set. Additionally, we selected another 2,900 ligands and pooled all their associated positive samples into the test set. The remaining datasets were split randomly into a training set and a validation set at a ratio of 10:1. The validation set was used to determine the hyperparameters, and the best model was evaluated on the test set. Under this experimental design, we finally established a ChEMBL set containing 217,732 samples in the training set, 24,193 samples in the validation set and 10,199 in the test set.

### External evaluation on external datasets

All the baseline models and TransformerCPI 2.0 were tested on the external test set, GPCR set and kinase set from our previous work. In addition to the GPCR and kinase set, we designed a large external set. This external set contains compounds that were not previously observed and 1,192 new protein targets that were not included in the training set. The total number of CPI pairs is 342,447, and the ratio of positive samples to negative samples is 1:3. This external set can evaluate the generalization ability of TransformerCPI2.0 and baseline models to the new compounds and new proteins. Another time-split dataset named the ChEMBL27 dataset contains compounds that were not previously observed and 637 new protein targets that are not included in the training set, and all the data were collected from the ChEMBL_27 database. The total number of CPI pairs is 92,919, and the ratio of positive samples to negative samples is 1:1.

### Baseline models

All the baseline models, including TransformerCPI, CPI-GNN, GraphDTA(GAT-GCN), and GCN, were trained on the ChEMBL dataset with their own hyperparameters. Only the learning rate, weight decay rate and dropout rate were subjected to a hyperparameter search.

### Drug resistance mutation analysis

Activity change score calculation. First, we input the wild-type protein and drug into TransformerCPI2.0 to calculate the original prediction score, denoted as *s*. Then, we mutated each amino acid of the protein sequence to all 20 amino acids (including itself) and calculated the prediction score *s*′. Finally, we defined the activity change score Δ*S* ∈ ℝ^*l*×20^ as

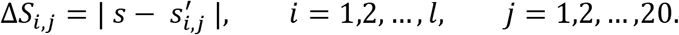

Here, *i* corresponds to the position of the protein sequence, *l* corresponds to the length of the protein sequence, and *j* corresponds to 20 types of amino acids. Since an amino acid mutation will increase or decrease the prediction score *s*′ and TransformerCPI2.0 may not be able to predict the trend of activity changes correctly, we calculated the absolute value of Δ*S* here. We analysed Δ*S* and found that TransformerCPI2.0 actually learns the key features of compound protein interactions because the pattern of Δ*S* revealed by heatmap analysis is consistent with that of drug resistance mutation.

### Relative activity change score

After calculating the activity change score, we can evaluate whether a mutation at a specific position plays an important role in compound protein interactions. However, Δ*S* cannot quantify the contribution of each position on a protein sequence and rank the most important sites from the whole sequence. To quantify the contribution of each amino acid site to drug-protein interactions, we first computed the average score of each position Δ^*S̅*^ ∈ ℝ^*l*^ as

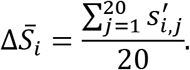

Furthermore, the value of Δ^*S̅*^ was normalized to between 0∼1, and the relative activity change score Δ*R* ∈ ℝ^*l*^ was defined as

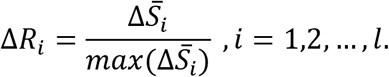

The relative activity change score Δ*R* can then characterize the contribution of each amino acid position to TransformerCPI2.0 prediction and help researchers discover novel and potential drug resistance mutation sites. On the other hand, Δ*R* can reflect the compound–protein interactions in TransformerCPI2.0. An amino acid site whose contribution to compound–protein binding is large can be revealed quantitatively by the relative activity change score Δ*R*. Finally, we used the activity change score Δ*S* to plot a heatmap to study the concrete pattern of drug resistance mutations and the relative activity change score Δ*R* to rank the most important sites for compound protein binding.

### Analysis of the substitution effect of the trifluoromethyl group

The replacement of methyl (Me or −CH3) with trifluoromethyl (TFM or −CF3) is frequently employed in compound optimization. However, the exact effect of −CH3/–CF3 substitution on bioactivity is still controversial. To further investigate whether TransformerCPI2.0 captures the key features of the compound and comprehensively understands compound–protein interaction, we employed TransformerCPI2.0 to predict the substitution effect of the trifluoromethyl group. We utilized a previously reported dataset^39^ containing 18,217 pairs of compounds and corresponding bioactivity data with the only difference being that −CH3 is substituted by −CF3 to study this problem.

### Dataset analysis

The statistical results showed that the replacement of −CH3 with −CF3 does not improve bioactivity on average^39^. However, in 15.73% of cases, substituting −CF3 for −CH3 increased or decreased the biological activity by at least an order of magnitude. Therefore, we focused on these samples to study the prediction capacity of TransformerCPI2.0. The actual bioactivity change Δ*pAct* was defined as

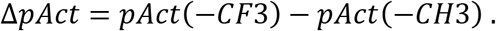

### Activity change score

First, we used TransformerCPI2.0 to calculate the prediction scores of compounds with trifluoromethyl substituents and denoted this score as *score*_−*CF*3_. Then, we calculated the prediction scores of compounds with methyl substituents and denoted this score as *score*_−*CH*3_. The activity change score Δ*s*_*c*_ was defined as

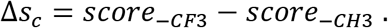

Considering that the distributions of Δ*pAct* and Δ*s*_*c*_ were different from each other, we defined a correct prediction by a model as Δ*pAct* and Δ*s*_*c*_ sharing the same sign. In other words, when the activity change trend predicted by the model matches the actual bioactivity change, the prediction is considered correct. After defining the evaluation metrics, we analysed the performance of TransformerCPI2.0 on the whole dataset. Furthermore, we investigated the prediction performance on the cases where the corresponding biological activities increased or decreased by at least three orders of magnitude because these cases are relevant to the activity cliff phenomenon in medicinal chemistry. Finally, we selected two cases that were not observed in the training set to show the power of TransformerCPI2.0.

### Comparison with structure-based docking tools

TransformerCPI2.0 is designed to predict compound–protein interactions considering only protein sequence information; however, evaluating the predictive power of TransformerCPI2.0 on those targets whose crystal structures remain unknown is not practical. Although we can test the power of TransformerCPI2.0 in specific lead compound screening projects, we compared TransformerCPI2.0 with structure-based docking tools to verify its ability to enrich positive molecules among compound libraries.

### Docking dataset

We used two benchmark datasets, DUD-E and DEKOIS 2.0, to compare TransformerCPI2.0 and other docking tools. DUD-E is a widely used benchmark dataset for docking tool and scoring function comparisons. DUD-E includes a total of 102 proteins with 22886 clustered ligands drawn from ChEMBL, each with 50 property-matched decoys drawn from ZINC. DEKOIS2.0 is often used as a complementary benchmark for further validation. DEKOIS2.0 contains 81 structurally diverse targets. Each target has 40 active compounds extracted from BindingDB and 1200 decoys generated from ZINC. DUD-E and DEKOIS 2.0 both serve as external test sets for TransformerCPI2.0, which means that TransformerCPI2.0 was not trained or fine-tuned on these datasets.

### Docking tools

Four docking programs, Glide SP (version 8.2)^65^, GOLD (version 2019)^66^ with Piecewise Linear Potential (CHEMPLP) SF, LeDock (version 1.0)^67^ and AutoDock Vina^68^, showed high docking power and computational efficiency and thus were chosen as the baseline models. We focused on the enrichment factor (EF0.5%, EF1%, EF5%) to evaluate different models, and the results of Glide SP, GOLD and LeDock on the two benchmark sets were extracted from Chao *et al*.^69^. In addition to the above four docking tools, many machine learning scoring functions have been tested on DUD-E and DEKOIS 2.0^69^; thus, we compared these machine learning scoring functions. We used the whole protein sequence as the default input of TransformerCPI2.0 and calculated the average EF0.5%, EF1% and EF5% of DUD-E and DEKOIS 2.0.

### Virtual screening of SPOP

First, after filtering non-drug-like compounds from the ChemDiv Library (San Diego, CA, USA), which contains approximately 1.6 million in-stock compounds, TransformerCPI2.0 was applied to score the compounds, and the top 35,000 molecules (∼top 2%, ensuring compound diversity) were selected by screening. Second, we filtered panassay interference compounds (PAINS) and clustered these molecules automatically based on their extended-connectivity fingerprints (ECFP), obtaining approximately 800 clusters. Third, we filtered these compounds by the Lipinski rules and manually selected representative compounds from each cluster. Finally, a total of 82 candidates were purchased for further experimental evaluation.

### Virtual screening of RNF130

First, after filtering non-drug-like compounds from the Chemspace Library (Monmouth Junction, NJ 08852, USA), which contains approximately 2 million in-stock compounds, TransformerCPI2.0 was applied to score the compounds, and the top 10,000 molecules (∼top 0.5%, ensuring compound diversity) were selected. Second, we filtered panassay interference compounds (PAINS) and clustered these molecules automatically based on their extended-connectivity fingerprints (ECFP), obtaining approximately 200 clusters. Third, we filtered these compounds by the Lipinski rules and manually selected representative compounds from each cluster. Finally, a total of 87 candidates were purchased for further experimental evaluation.

### Target identification of PPIs

We collected potential proteins from the DrugBank database and selected proteins that already have active modulators. Then, TransformerCPI2.0 was applied to score proteins against four classical PPIs (omeprazole, rabeprazole, lansoprazole and pantoprazole), and the results were sorted by predicted interaction probability. Next, we analysed the top 20 proteins by evaluating their novelty, importance and feasibility, and finally chose ARF1 for experimental validation.

### Compounds

SPOP inhibitors were purchased from ChemDiv Library (San Diego, CA, USA): 221C7, Y502-3210; 231A10, 5282-0816; 231D8, 8017-3040. 230D7 was synthesised in our laboratory. RNF130 inhibitor was purchased from Chemspace Library (Monmouth Junction, NJ 08852, USA): iRNF130-63, CSC138461036. PPIs were purchased from MedChemExpress (Monmouth Junction, NJ, USA): rabeprazole, HY-B0656; lansoprazole, HY-13662; Omeprazole, HY-B0113; pantoprazole, HY-17507.

### Plasmid construction

Wild-type, truncated or mutant versions of the human proteins were used in this study: SPOP (UniProt accession number: O43791-1), PTEN (UniProt accession number: P60484-1), DUSP7 (UniProt accession number: Q16829-1), RNF130 (UniProt accession number: Q86XS8-1), ARF1 (UniProt accession number: P84077-1) and ARNO (UniProt accession number: Q99418-1). For plasmid construction, SPOP^WT^ and SPOP^cyto^ (residues 1-366) were subcloned into the pcDNA 3.1 vector with a Flag-tag, SPOP^MATH^ (residues 28-166) and ARNO^Sec7^ (residues 50-250) was subcloned into the pGEX 6p-1 vector with a GST-tag, PTEN and DUSP7 were subcloned into the pcDNA 3.1 vector with a Myc-tag, RNF130 (residues 1-304) was subcloned into the pcDNA3.1 vector with a C-terminal His_8_-Flag-tag. N-terminally truncated human Δ17ARF1 and C159A-mutant Δ17ARF1 were subcloned into pProEX HTb with a His_6_-tag. All of the above plasmids were synthesized by Sangon Biotech (Shanghai) Co., Ltd. pCMV-HA-Ub plasmid (CAT#. kl-zl-0513) was purchased from Shanghai Kelei Biological Technology Co., Ltd.

### Recombinant protein expression and purification

For expression of SPOP^MATH^, GST-tagged SPOP^MATH^ plasmid was transformed into BL21-CodonPlus (DE3)-RIPL Cells (CAT#. EC1007, Shanghai Weidi Biotechnology Co., Ltd.), and then the cells were grown in lysogeny broth (LB) medium and induced by isopropyl β-D-1-thiogalactopyranoside (IPTG) at a final concentration of 0.5 mM at 16 °C overnight. Cells were harvested and lysed in solution (20 mM HEPES pH 7.4, 200 mM NaCl, 1 mM dithiothreitol) by sonication and then centrifuged at 16,000 rpm for 1 hour at 4 °C. The supernatants were filtered by 0.22 μM syringe filters and purified on GST Trap columns (GE Healthcare) by elution with 10 mM glutathione. The eluted components were loaded onto desalting columns (GE Healthcare) to remove glutathione and incubated with PreScission Protease for 6-8 hours at 4 °C. The components were reloaded onto GST Trap columns to remove the GST tags and further purified by a Superdex 75 10/300 GL column. The purified SPOP^MATH^ protein was concentrated and stored in buffer (20 mM HEPES pH 7.4, 200 mM NaCl) at -80 °C.

RNF130 protein was expressed in Expi-293F (Invitrogen) using Expifectamine transfection reagent according to the manufacturer’s instructions. Cells were collected 3 days after transfection. Proteins were first captured by Ni^2+^-Sepharose 6 Fast Flow resin (GE healthcare) and then further purified by gel filtration chromatography with a Superdex S200 column (GE Healthcare). The purified protein was concentrated and stored in buffer (20 mM HEPES pH 7.5, 150 mM NaCl and 1 mM TCEP) at -80 ℃. For expression of Δ17ARF1 and Δ17ARF1^C159A^, the His-tagged recombinant plasmid was transformed into BL21-CodonPlus (DE3)-RIPL cells, and then the cells were grown in LB medium and induced by IPTG at a final concentration of 0.1 mM at 25 °C for 6 hours. The cells were harvested and lysed in solution (20 mM Tris, 100 mM NaCl, 5 mM MgCl_2_, 10 mM imidazole, pH 8.0) by sonication and then centrifuged at 16,000 rpm for 1 hour at 4 °C. The supernatants were filtered by 0.22 μM syringe filters and purified on HiTrap column (GE Healthcare) by elution with 300 mM imidazole. Protein sample was then purified by a Superdex 75 10/300 GL column. Finally, the purified Δ17ARF1 and Δ17ARF1^C159A^ proteins were concentrated and stored in buffer (20 mM Tris, 100 mM NaCl, pH 8.0) at -80 °C.

The purification process ARNO^Sec7^ protein was the same as that of the SPOP^MATH^ protein except the cells were induced by 0.1 mM IPTG at 37 °C for 3 hours.

### Fluorescence polarization (FP)

Fluorescence polarization experiments were conducted in a 384-well black plate (Corning, 3575) using a 42 μL reaction system. FITC-labelled SPOP substrate puc_SBC1 (FITC-LACDEVTSTTSSSTA)^70,71^ (synthesized by GL Biochem (Shanghai) Ltd) was used for the probe. Then, 20 μL of reaction buffer (20 mM HEPES, pH 7.4) containing 200 nM SPOP^MATH^ protein was incubated with 2 μL of compound for 30 minutes at room temperature, and 20 μL of reaction buffer containing 200 nM probe was added. Fluorescence polarization (mP) signals were measured by a fluorescence mode (excitation filter 480 nm, emission filter 535 nm) in Spark microplate reader (Tecan).

### Protein thermal shift (PTS)

The Bio–Rad CFX96 RealTime PCR Detection System was utilized to monitor the thermal stability of ARF1 and SPOP^MATH^ protein. PTS experiments were performed in a 96-well PCR plate (DN Biotech (Hong Kong) Co., Ltd.) with a 20 μL reaction system. A total of 20 μL of reaction buffer containing protein (5 μM for SPOP^MATH^ or 2.5 μM for ARF1), 5 × SYPRO Orange Protein Gel Stain (Sigma, S5692) and indicated concentration of compound. The signals of all reaction systems were continuously monitored and recorded from 25 °C to 90 °C for approximately 45 minutes. The T_m_ values of SPOP^MATH^ and ARF1 were measured using CFX manager software version 3.1.

### Nuclear magnetic resonance (NMR)

NMR spectroscopy experiments were performed using a 600 MHz spectrometer (AVANCE III, Bruker) to validate protein– ligand interactions. In Carr-Purcell-Meiboom-Gill (CPMG) and saturation transfer difference (STD) NMR experiments, compound was dissolved to a final concentration of 200 μM in a solution of PBS formed with D_2_O containing 5 μM SPOP^MATH^ protein and 5% DMSO-d^6^.

### Mass spectrometry analysis

The experiment was performed on the mass spectrometry service platform of Shanghai Institute of Materia Medica, Chinese Academy of Sciences. The protein (100 μM) was incubated with compounds (1 mM) or solvent control overnight at 4 °C, and then the protein molecular weights were determined by Q Exactive (Thermo) and 6545 XT (Aglient) mass spectrometer. For compound binding site identification, the proteins were digested with trypsin (10 ng/μL) at 37 °C for 17 h. The next day, after centrifugation, the supernatant was lyophilized, desalted, and lyophilized again, followed by the addition of 0.1% FA solution to dissolve peptide lyophilized powder. After centrifugation, the supernatant was detected by mass spectrometry (Q-Exactive). The MS data was analysed via software MaxQuant (version 1.6.5.0). The false discovery rate (FDR) for peptides and proteins was controlled <1% by Andromeda search engine.

### Surface plasmon resonance (SPR) for RNF130

The SPR binding assay was performed using a Biacore T200 instrument (GE Healthcare). The purified RNF130 protein was covalently immobilized onto a CM5 sensor chip (Cytiva) by a standard amine-coupling procedure in 10 mM sodium acetate (pH 4.5) with running buffer HBS (50 mM HEPES pH 7.4, 150 mM NaCl). iRNF130-63 was serially diluted and injected onto the sensor chip at a flow rate of 30 μL/min for 120 s (contact phase), followed by 120 s of buffer flow (dissociation phase). The equilibrium dissociation constant (K_D_) value was derived using Biacore T200 Evaluation software (version 1.0, GE Healthcare).

### Guanine nucleotide exchange assay for ARF1

First, with the participation of EDTA (a metal chelating agent capable of chelating magnesium ions, which are critical for the binding of GDP/GTP to ARF1), GDP was loaded onto the ARF1 protein by incubating ARF1 with a 20-fold molar concentration of GDP. Excess magnesium chloride was used to terminate the loading reaction, followed by the removal of excess GDP by a NAP-5 column to produce ARF1^GDP^ protein. Next, ARF1^GDP^ protein (20 μM) was mixed with compounds and Mant-GTP (10 μM) in reaction buffer and incubated in the dark for 15 minutes. Exchange reactions were initiated by the injection of ARNO^Sec7^ (1 μM).

### Cell lines

293T, MDA-MB-231, 4T-1 and CT26 cells were obtained from the American Type Culture Collection (ATCC). OS-RC-2 and Caki-2 cells were purchased from the National Biomedical Laboratory Cell Resource Bank. 786-O cells were kindly provided by the Cell Bank/Stem Cell Bank, Chinese Academy of Sciences. 293T and MDA-MB-231 cells were cultured in DMEM medium (BasalMedia, L110KJ) supplemented with 10% foetal bovine serum (FBS, Gibco, 10099141C) and 1% Penicillin-Streptomycin (PS, Gibco, 2321118). 786-O, OS-RC-2, 4T-1 and CT26 cells were cultured in RPMI-1640 medium (BasalMedia, L210KJ) supplemented with 10% FBS and 1% PS. Caki-2 cells were cultured in McCoy’s 5A medium (Gibco, 2193071) supplemented with 10% FBS and 1% PS. All cells were incubated at 37 °C under a 5% (v/v) CO_2_ atmosphere.

### G-LISA

A Cytoskeleton ARF1 G-LISA Activation Assay Kit (BK132) was used to detect cellular ARF1-GTP levels according to the manufacturer’s instructions.

### Western blot

Total proteins from cells were lysed in RIPA lysis buffer (Beyotime, P0013C) containing phosphatase inhibitor (Bimake, B15001) and protease inhibitor (Bimake, B14001) on ice. Cell lysates were centrifuged at 13,000 × g for 15 minutes at 4 °C. The BCA protein assay kit (Thermo Scientific, 23225) was used to quantify the protein concentration. Equal amounts of total proteins were separated by 10% SDS– PAGE and then transferred onto nitrocellulose membranes. The membranes were blocked with 5% skim milk in TBST for 1 hour at room temperature and then incubated with primary antibodies overnight at 4 °C. Then membranes were incubated with HRP-conjugated anti-rabbit antibody (secondary antibody, Promega, W4011) for 1 hour at room temperature. Finally, the immune complexes were detected with an ECL kit (Meilun, MA0186) and visualized as well as quantified using GenGnome XRQ NPC. The following primary antibodies were used: anti-Flag (Cell Signaling Technology, 14793), anti-myc (Cell Signaling Technology, 2278), anti-GAPDH (Cell Signaling Technology, 5174), anti-PTEN (Cell Signaling Technology, 9559), anti-HA (Cell Signaling Technology, 3724), anti-p-ERK^T202/Y204^ (Cell Signaling Technology, 4376), anti-p-AKT^Thr308^ (Cell Signaling Technology, 4056), anti-ERK (Cell Signaling Technology, 9102), anti-AKT (Cell Signaling Technology, 9272), anti-DUSP7 (ABGENT, AP8450a), anti-SPOP (Abcam, ab192233), anti-GST (Absin, abs830010) and anti-β-Tubulin (Cell Signaling Technology, 15115).

### *In vitro* GST pull-down

The plasmids (Myc-PTEN or Myc-DUSP7) were transiently transfected into 293T cells. After transfection for 24 hours, the cells were harvested and lysed in cell lysis buffer for Western and IP (Beyotime, P0013) containing protease inhibitor on ice. GST or GST-SPOP^MATH^ proteins bound to GST magnetic beads (GenScript, L00327) were incubated with the cell lysates (Myc-PTEN or Myc-DUSP7) in the presence of different doses of compound for 2 hours at room temperature. The beads were washed 3 times with PBST, and the precipitated proteins were eluted with 1 × SDS loading buffer (Beyotime, L00327) at 100 °C for 5 minutes and analysed by Western Blot.

### Cellular thermal shift assay (CETSA)

293T cells transfected with Flag-RNF130 for 48 hours were collected and lysed in 20 mM Tris pH 7.5, 150 mM NaCl and 1% Triton X-100. Then, 50 μM iRNF130-63 or DMSO was added to the supernatant and incubated at 25 °C for 30 minutes. After denaturing at various temperatures for 3 minutes on a temperature gradient PCR instrument (Eppendorf), the samples were centrifuged at 20000 × g for 30 minutes at 4 °C, and the supernatants were analysed by Western blot.

### Coimmunoprecipitation (Co-IP)

The plasmids (Flag-SPOP^cyto^, Myc-PTEN or Myc-DUSP7) were transiently cotransfected into 293T cells. After transfection for 24 hours, 293T cells were treated with different doses of compound for another 24 hours. The cells were harvested and lysed in cell lysis buffer for Western and IP containing protease inhibitor on ice. Approximately 80% of the total lysates were immunoprecipitated with anti-Flag-conjugated magnetic beads (Bimake, B26102) for 2 hours at room temperature, and other lysates were used as input. The magnetic beads were then washed 3 times with PBST, and the immunoprecipitated proteins were eluted with 1 × SDS loading buffer at 100 °C for 5 minutes. The IP and lysate samples were analysed by Western Blot.

### *In vivo* ubiquitination

The plasmids (Myc-PTEN or Myc-DUSP7, Flag-SPOP^cyto^, HA-Ub) were transiently cotransfected into 293T cells. After transfection for 24 hours, 293T cells were treated with different doses of compound for 24 hours. The cells were then treated with 10 μM protease inhibitor MG132 (MedChemExpress (Monmouth Junction, NJ, USA), HY-13259) for another 4 hours before harvesting. Next, the cells were lysed in cell lysis buffer for Western and IP containing protease inhibitor on ice. Approximately 80% of the total lysates were immunoprecipitated with anti-Myc-conjugated magnetic beads (Bimake, B26302) for 2 hours at room temperature, and the other lysates were used as input. The magnetic beads were then washed 3 times with PBST, and the immunoprecipitated proteins were eluted with 1 × SDS loading buffer at 100 °C for 5 minutes. The ubiquitination levels were detected using a Western Blot assay.

### Cell proliferation

Cells were seeded in 96-well plates and incubated with serially diluted compounds for 72 hours. Cell viability was determined using the CellTiter-Glo® Luminescent Cell Viability Assay kit (Promega, G7573) following the manufacturer’s instructions. IC_50_ values were determined by nonlinear regression (curve fit) using a variable slope (four parameters) in GraphPad Prism (9.0).

### Animals

All procedures performed on animals were in accordance with regulations and established guidelines and were reviewed and approved by the Institutional Animal Care and Use Committee at the Shanghai Institute of Materia Medica, Chinese Academy of Sciences (IACUC Issue NO. 2022-01-JHL-27 for NSG mice; IACUC Issue NO. 2021-03-JHL-22 for BALB/c mice). NSG mice were obtained from Shanghai Model Organisms Center, Inc; BALB/c mice were obtained from Beijing Huafukang Biotechnology Co. Ltd (Beijing, China). Six-to eight-week-old mice were used for the studies and were maintained with free access to pellet food and water in plastic cages ate 21 ± 2℃ and kept on a 12 hours light/dark cycle.

### Pharmacokinetics

The pharmacokinetic profiles of compound 230D7 were determined in male BALB/c mice. The test compound 230D7 was dissolved in solution containing DMSO, PEG400, PBS (5/5/90, v/v) and administered via intraperitoneal administration (i.p.) at 10 mg/kg. Serial blood samples were collected at 0.25, 0.5, 1, 2, 4, 8, 24 hours after dosing and centrifuged to obtain the plasma fraction. A 10 μL aliquot of plasma was deproteinized with 100 μL acetonitrile/methanol (1/1, v/v) containing internal standard. After centrifugation, the supernatant was diluted with a certain proportion of acetonitrile/water (1/1, v/v), mixed and centrifuged at 4,000 rpm for 10 minutes. Finally, the aliquots of the diluted supernatant were injected into LC-MS/MS system.

### Acute toxicity

BALB/c mice were used to evaluate the toxicity of compound 230D7. The mice were randomly divided into 3 groups (n = 3) and treated with different doses of compound 230D7 (0, 50, 100 mg/kg) by intraperitoneal administration daily for a week. The body weights of mice were measured every day and the significant organs (heart, kidney, lung, liver and spleen) were harvested, weighted and used for histological analysis at the last day.

### H&E staining

For histological analysis of BALB/c mice in 230D7-treated or vehicle control groups, staining for H&E staining were performed using standard histological techniques. Isolated organ tissues were fixed in neutral paraformaldehyde for 24 h and embedded in paraffin wax. H&E staining of these tissue examples were carried out by Servicebio company.

### 786-O cells xenograft tumour growth

NSG mice were used to evaluate the pharmacodynamics of 230D7. The 786-O cells xenograft tumour model was established by the subcutaneous injection of 786-O cells (1 × 10^7^) into the NSG mice. When the tumour reached the volume of approximately 100 mm^3^, the mice were randomly divided into three groups (n = 7) and intraperitoneally treated with different dosages of 230D7 (0, 25, 50 mg/kg, 230D7 was synthesised in our laboratory) in solution containing DMSO, PBS (5/90, v/v) once a day for 16 days. Body weight and tumour size were measured every two or three days, and tumour volume was calculated using the formula: V = (L × W^2^)/2 (L, length; W, width). At the end of the experiment, the mice were euthanized and the tumours were harvested for Western Blot and other studies.

### CT26 cells transplanted tumour growth

BALB/c mice were used to evaluate the pharmacodynamics of rabeprazole (purchased from MedChemExpress, HY-B0656). The CT26 cells transplanted tumour model was established by the subcutaneous injection of CT26 cells (2.5 × 10^5^) into mice. When the tumour reached the volume of approximately 100 mm^3^, the mice were randomly divided into two groups (n = 5) and intraperitoneally treated with rabeprazole (0, 40 mg/kg) in solution containing DMSO, PEG300, PBS, (1/10/89, v/v/v) once a day for 10 days. The tumour size was recorded using callipers, and tumour volume was calculated using the formula: V = (L × R^2^)/2. At the end of the experiment, the mice were euthanized and the tumours were harvested for immunohistochemistry and fluorescence-activated cell sorting.

### Nile red staining

CT26 cells were seeded into Lab-Tek^TM^ II Chamber Slide systems (Thermo) and incubated with rabeprazole or vehicle. Then, the cells were washed with PBS for 15 minutes, fixed with Immunol Staining Fix Solution (Beyotime, P0098) for 30 minutes and washed with PBS again, followed by treatment with Immunostaining Permeabilization Buffer with Triton X-100 (Beyotime, P0098) for 30 minutes and washing with PBS again. To stain the lipid droplets, the cells were incubated with Nile Red (2 µM) in the dark for 10-30 minutes and then washed with PBS before the nuclei were stained with Antifade Mounting Medium with DAPI (Beyotime, P0131). Fluorescence images were captured using an OLYMPUS IX73 fluorescence microscope.

### Immunohistochemistry (IHC)

The isolated CT26 tumour tissue was fixed with neutral paraformaldehyde, and subsequent staining of cell surface markers was performed by Servicebio Company.

### Fluorescence-activated cell sorting (FACS)

We analyzed the infiltration of immune cell subsets in CT26 transplanted tumour tissue by fluorescence-activated cell sorting (FACS) analysis. After the mice were euthanized, the tumour tissues were stripped and cut into pieces, then digested at 37 ℃ for 60 minutes with tumour tissue digestive buffer (0.1% collagenase, 0.001% hyaluronidase, 0.002% DNA enzyme, 120 μM CaCl_2_ and 120 μM MgCl_2_ in RPMI1640 medium). The digested tumour tissues were filtered with 200 mesh gauze, followed by the lysis of red blood cells with ammonium chloride solution, and filtered again to obtain single-cell suspensions in PBS. For discriminating the living and dead cells, single-cell suspensions were stained on ice with Fixable Viablity Stain 700 (BD Horizan, 564997) for 10 min and then terminated with cell staining buffer (PBS containing 2% FBS). Fc receptors on the cell surface are blocked by anti-Mouse CD16/32 antibody (10 minutes on ice). Then, appropriately conjugated fluorescent primary antibodies were added to stain cell surface markers. Finally, cells were suspended with cell staining buffer for flow cytometry analysis using Beckman CytoFelx. The data were analyzed by Flowjo sofrware.

### Statistical analysis

GraphPad Prism 9.0 software was used to perform statistical analysis. Differences of quantitative data between groups were calculated using 2-tailed unpaired t-test. The significance level was set as ^*^*P* < 0.05, ^**^*P* < 0.01, ^***^*P* < 0.001, ^****^*P* < 0.0001.

### Synthesis of 230D7

Unless otherwise specified, all reagents and solvents were purchased from commercial sources and used without further purification. ^1^HNMR spectra was recorded on Mercury-600 spectrometers at room temperature. Chemical shifts are referenced to the residual solvent peak and reported in ppm (δ scale), and all coupling constant (*J*) values are given in Hz. ESI-LRMS data were measured on Thermo Exactive Orbitrap plus spectrometer. Flash column chromatography was performed on Flash 300 Isolera one. ^1^H NMR and ESI-LRMS data were reported on Extended Data Fig. 9.

Scheme I. Synthetic Route to Compound **230D7**

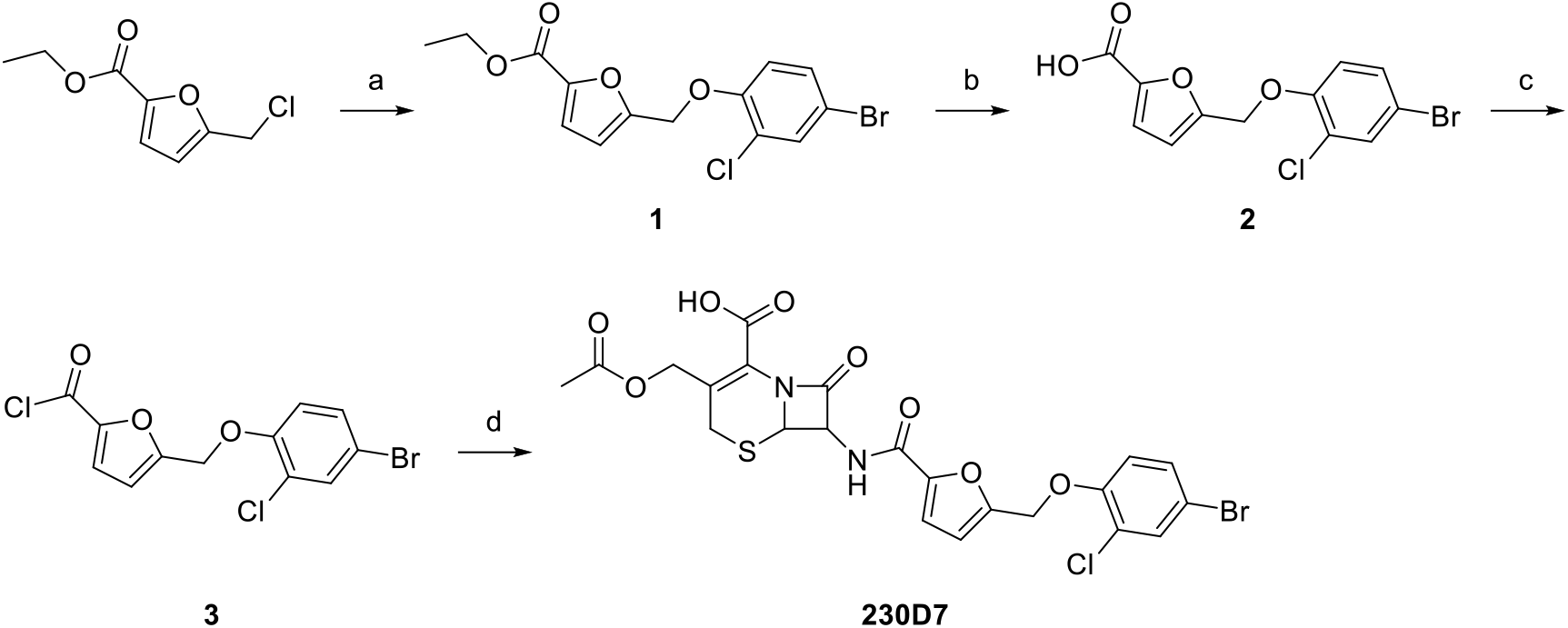

Reagents and conditions: (a) 4-bromo-2-chlorophenol, K_2_CO_3_, DMF, 60 °C, 5 h; (b) NaOH, MeOH, H_2_O, 50 °C, 2 h; (c) SOCl_2_, r.f., 2 h; (d) β-lactam amine, NaHCO_3_, acetone, H_2_O, r.t., 4 h.

#### ethyl 5-((4-bromo-2-chlorophenoxy)methyl)furan-2-carboxylate (1)

To a solution of 4-bromo-2-chlorophenol (1.0 g, 4.9 mmol) and K_2_CO_3_ (1.1 g, 8.0 mmol) was added 7 mL of DMF at 60 °C for 5 h. After reaction completion, H_2_O (50 mL) was added to the mixture followed by extraction with EtOAc (3 × 50 mL). The organic phases were collected and evaporated. The crude product was purified by column chromatography and eluted with 20% ethyl acetate in petroleum ether as a solvent to obtain a white solid product **1** (1.2 g) in 63% yield.

#### 5-((4-bromo-2-chlorophenoxy)methyl)furan-2-carboxylic acid (2)

To a solution of **1** (1.2 g, 3.4 mmol) and NaOH (0.48 g, 10.2 mmol) was added 15 mL of MeOH and 15 mL of H_2_O at 50 °C for 5 h. After reaction completion, MeOH was removed under reduced pressure. In the reaction mixture, 1.0 M aqueous HCl (5.0 mL) was added and pH value was adjusted to 4 to precipitate 1.09 g product **2** without further purification in 98% yield.

#### 5-((4-bromo-2-chlorophenoxy)methyl)furan-2-carbonyl chloride (3)

To a solution of **2** (0.20g, 0.61 mmol) in SOCl_2_ (3.0 mL) was stirred and heated at reflux for 2 h. After reaction completion, SOCl_2_ was removed under reduced pressure to obtain 0.21 g desired product **3** without further purification for next step in 100% yield.

#### 3-(acetoxymethyl)-7-(5-((4-bromo-2-chlorophenoxy)methyl)furan-2-carboxamido)-8-oxo-5-thia-1-azabicyclo[4.2.0]oct-2-ene-2-carboxylic acid (230D7)

3-(acetoxymethyl)-7-amino-8-oxo-5-thia-1-azabicyclo[4.2.0]oct-2-ene-2-carboxylic acid (0.21 g, 0.76mmol) was dissolved in 20 mL sat. aqueous NaHCO3 and 7 mL acetone at 0 °C. To the stirring solution, **3** (0.32 g, 0.91 mmol) in 5 mL acetone was added to reaction mixture dropwise in 15 min. After ice water bath was removed, reaction was allowed to stir at room temperature for another 4 h. After reaction completion, pH value was adjusted to 4 with 1.0 M aqueous HCl. Subsequently, the mixture was filtered to collect precipitate as crude product. The crude product was purified by column chromatography and eluted with 5% MeOH in DCM as a solvent to obtain 0.30 g white solid product (**230D7**) in 68% yield. ^1^H NMR (600 MHz, DMSO-*d*_6_) δ 13.71 (s, 1H), 9.35 (d, *J* = 8.0 Hz, 1H), 7.72 (d, *J* = 3.0 Hz, 1H), 7.45 (dd, *J* = 9.0, 3.0 Hz, 1H), 7.38 (d, *J* = 3.0 Hz, 1H), 7.33 (d, *J* = 9.0 Hz, 1H), 6.78 (d, *J* = 3.0 Hz, 1H), 5.82 (dd, *J* = 8.0, 5.0Hz, 1H), 5.24 (s, 2H), 5.18 (d, *J* = 5.0 Hz, 1H), 5.00 (d, *J* = 13.0 Hz, 1H), 4.71 (d, *J* = 13.0 Hz, 1H), 3.65 (d, *J* = 18.0 Hz, 1H), 3.51 (d, *J* = 18.0 Hz, 1H), 2.04 (s, 3H), ESI-MS: calcd for C_22_H_18_BrClN_2_O_8_S [M-H]^-^: 584.9, found: 584.9.

### Data availability

ChEMBL database is available at https://www.ebi.ac.uk/chembl/, DUDE dataset is available at http://dude.docking.org/, DEKOIS2.0 dataset is available at http://www.pharmchem.uni-tuebingen.de/dekois/, and DrugBank dataset is available at https://go.drugbank.com/. The commercial Chemspace Library is available at https://chem-space.com/, and ChemDiv Library is available at https://www.chemdiv.com/. All data are available in the main text or the supplementary materials.

### Code availability

Source code of TransformerCPI is available at https://github.com/lifanchen-simm/transformerCPI. The details of upgrading TransformerCPI to TransformerCPI2.0\ are provided in the Method section.

## Competing interests

The Shanghai Institute of Materia Medica has applied for Chinese patents that covers SPOP compounds.

## Acknowledgements

We gratefully acknowledge financial support from Lingang Laboratory (LG202102-01-02 to M.Z. and LG-QS-202204-01 to S.Z.), the National Natural Science Foundation of China (81903639 to S.Z.), the Shanghai Municipal Science and Technology Major Project and the Natural Science Foundation of Shanghai (22ZR1474300 to S.Z.).

## Author Contributions

L.C., Z.F., J.C. and R.Y. designed and performed the experiments, prepared the figures, and wrote the manuscript; L.C. designed TransformerCPI2.0 and conducted computational work; T.Y. conducted computational analysis; Z.F., C.Z., Z.C., R.C. and R.Y. contributed to the biological experiments on SPOP inhibitors; H.G., C.Z. and X.H. contributed to the synthesis of SPOP inhibitors; Y.D. and N.Z. contributed to nuclear magnetic resonance experiments on SPOP inhibitors; R.Y. contributed to the biological experiments on RNF130; and J.C., Y.Z., K.Z. and R.Y. contributed to the biological experiments on ARF1 inhibitors. M.Z., S.Z. and H.J. initiated, designed and supervised this study.

---

**Extended Data Fig. 1.**
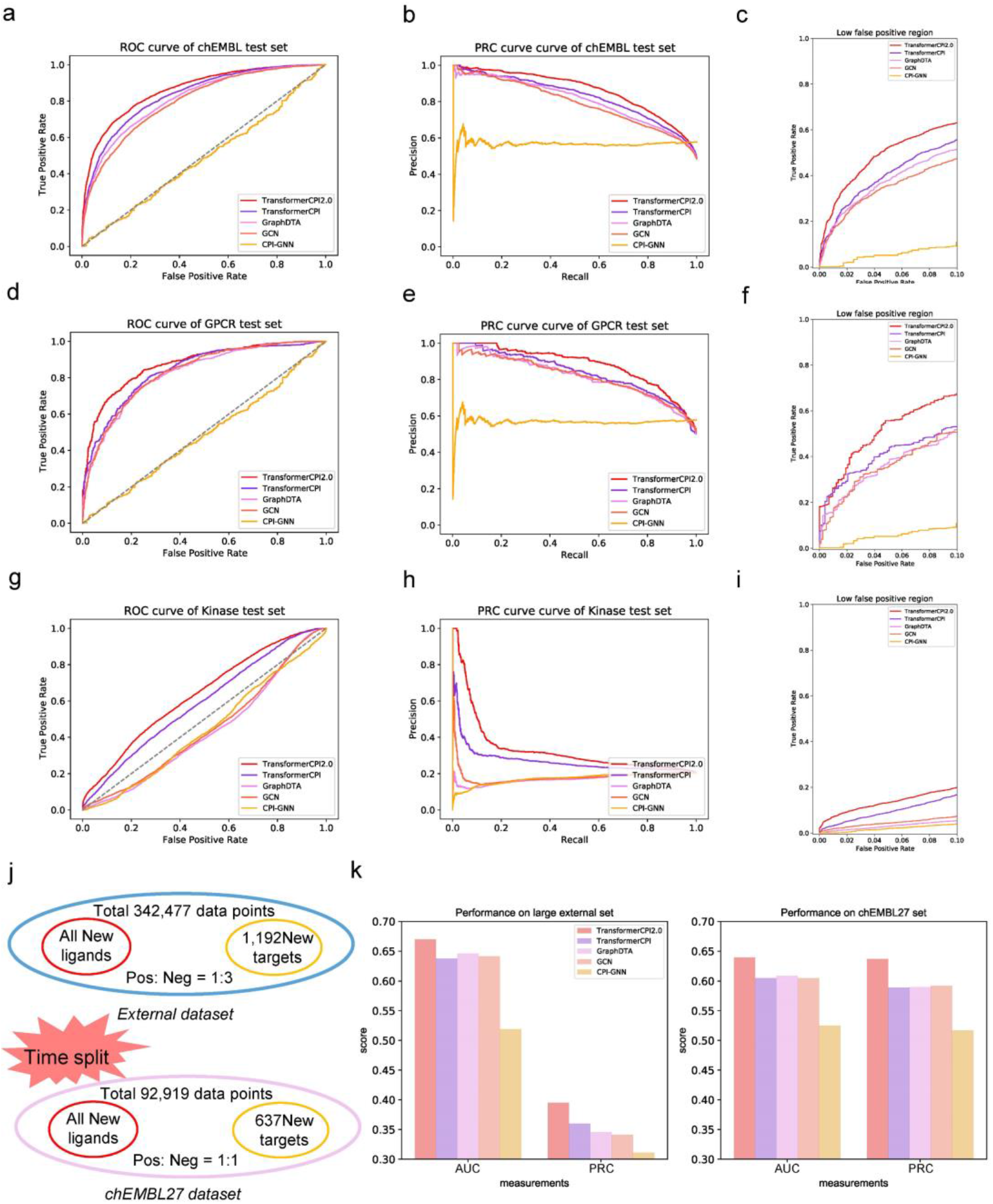
TransformerCPI 2.0 predicts compound–protein interaction without protein structure. **a,** AUC curves of TransformerCPI2.0 and baseline models on the chEMBL set. **b**, PRC curves of TransformerCPI2.0 and baseline models on the chEMBL set. **c**, ROC curves in low-false-positive region. **d,** AUC curves of TransformerCPI2.0 and baseline models on the GPCR set. **e**, PRC curves of TransformerCPI2.0 and baseline models on the GPCR set. **f**, ROC curves in a low-false-positive region. **g**, AUC curves of TransformerCPI2.0 and baseline models on the Kinase set. **h**, PRC curves of TransformerCPI2.0 and baseline models on the Kinase set. **i,** ROC curves in a low-false-positive region. **j**, The arrangement of the external dataset and chEMBL27 dataset. **k**, The performance of TransformerCPI2.0 and baseline models on the external set and chEMBL27 set.

**Extended Data Fig. 2.**
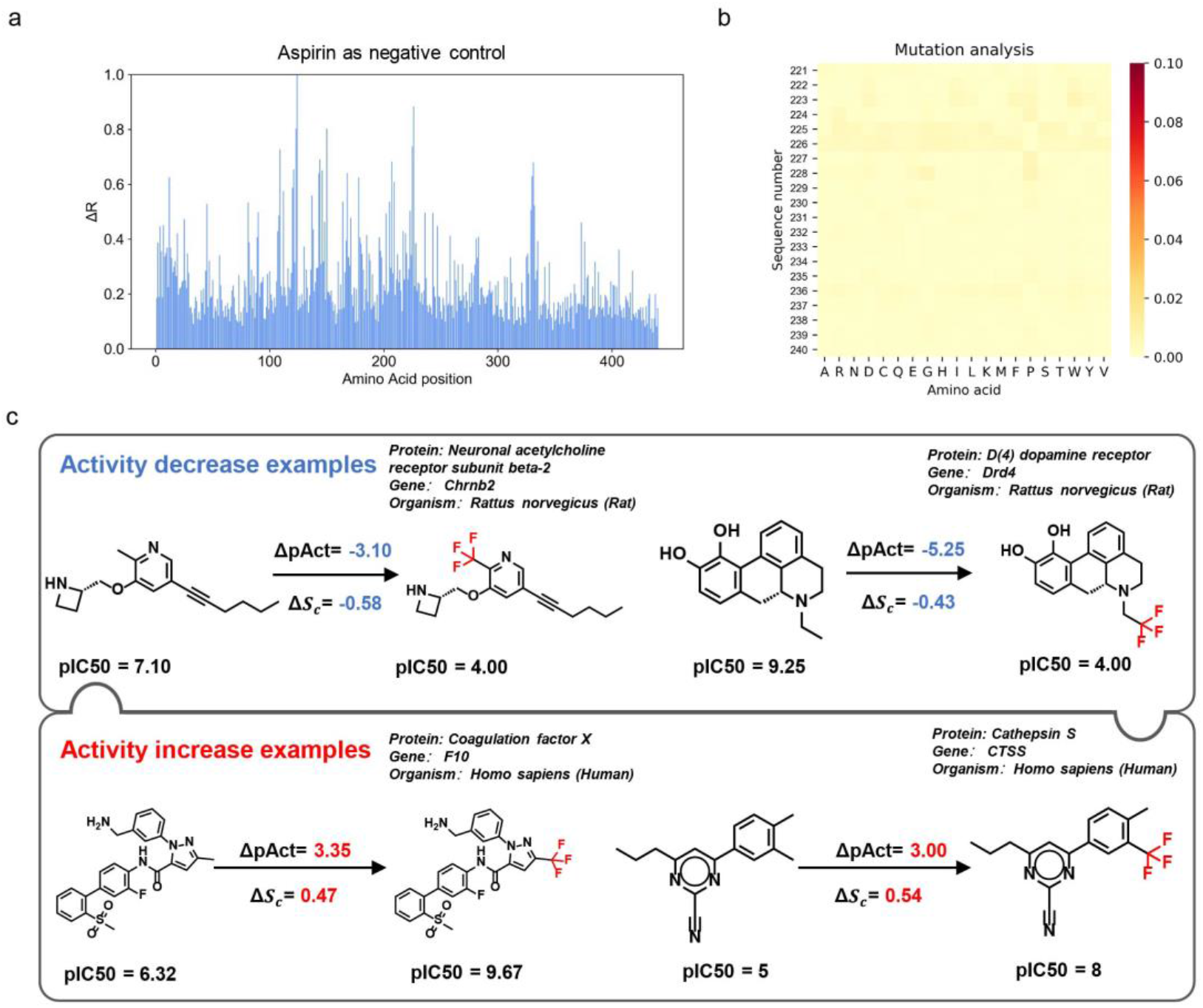
Supplementary data from drug resistance mutation analysis and substitution effect analysis of the trifluoromethyl group. **a,** Relative activity change score (Δ*R*) calculated by TransformerCPI 2.0 for each amino acid position. The pink boxes mark the high Δ*R* regions, which are plotted in detail on the right. **b**, The heatmap plots the activity change score of positions 221 to 240, where each position is mutated to the 20 different amino acids (including itself). The darker colour represents the higher activity change score caused by the mutation. **c**, Additional two activity decrease examples and activity increase examples are shown. The predictions of TransformerCPI2.0 are consistent with the ground truth for proteins and compounds not present in the training set.

**Extended Data Fig. 3.**
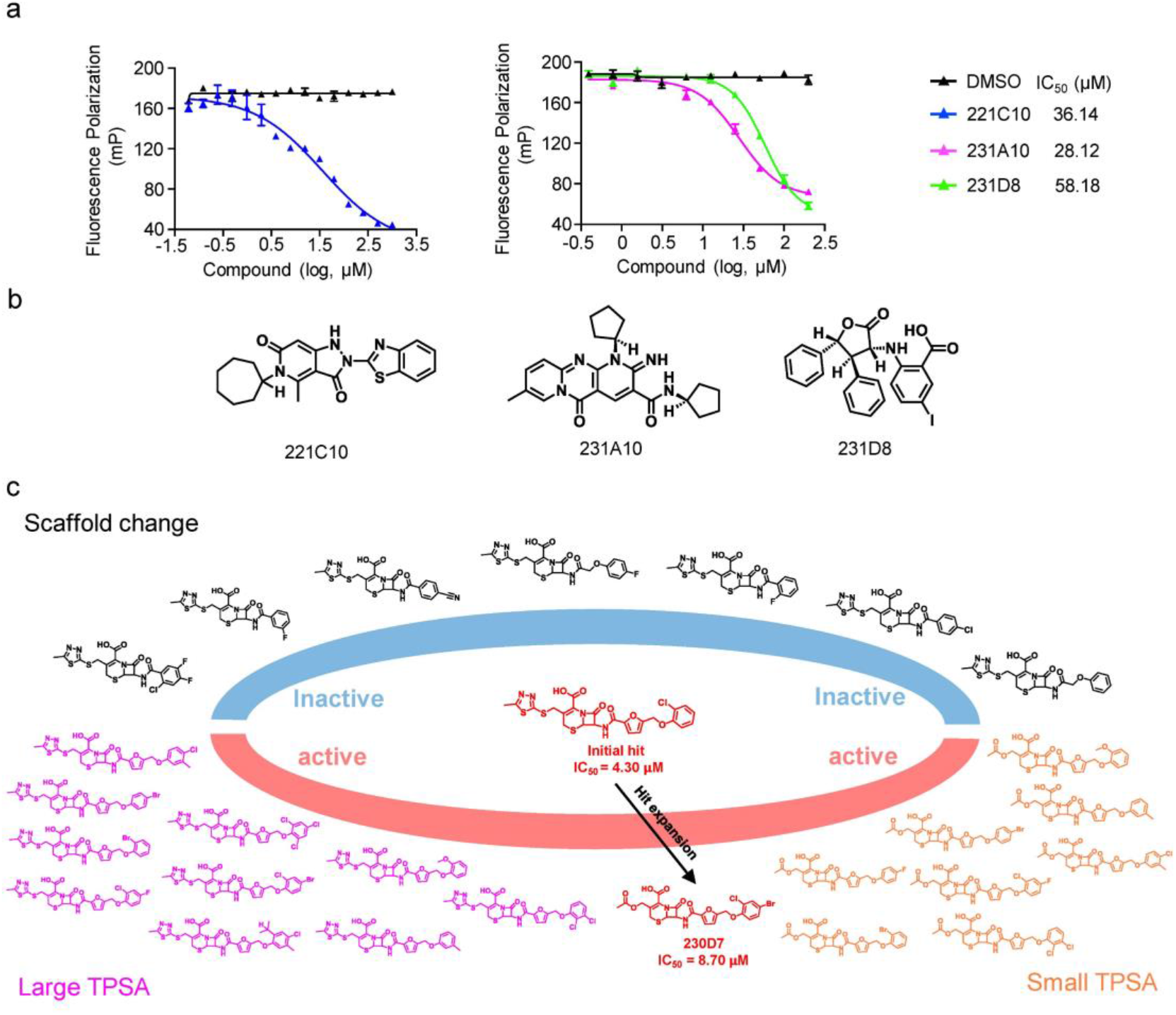
Supplementary data for Fig. 3. **a**, The FP assay of three initial hits predicted by TransformerCPI2.0. **b**, Chemical structures of the three initial hits predicted by TransformerCPI2.0. **c**, A similarity search of 221C7 was conducted, and 26 compounds were purchased, 19 of which were active in the FP assay.

**Extended Data Fig. 4.**
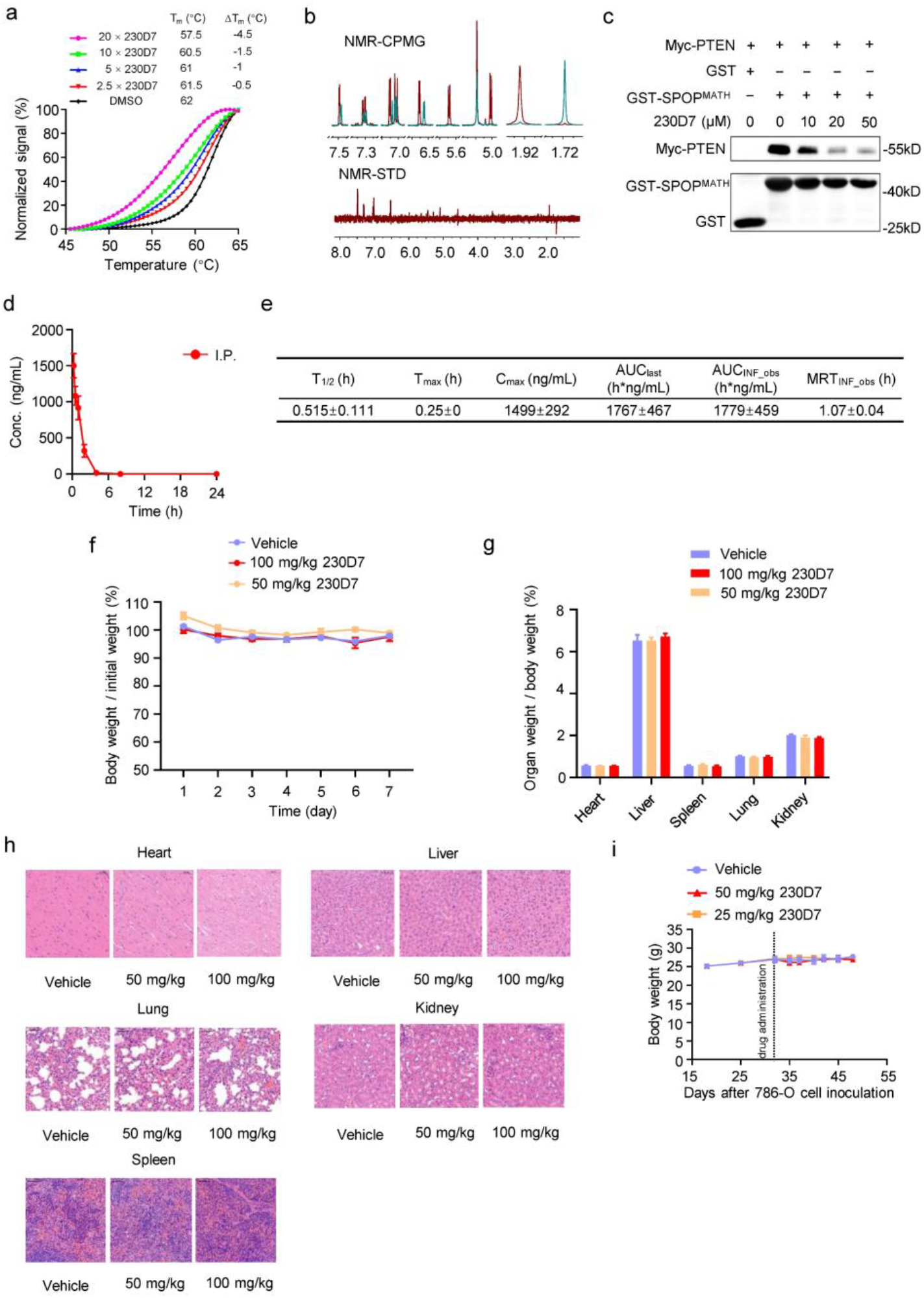
Supplementary data for Fig. 4. **a,** Thermostability of SPOP^MATH^ treated with different concentrations of 230D7. The thermal stability of SPOP^MATH^ was quantified by the ΔT_m_. **b**, NMR measurement of direct binding between 230D7 and SPOP^MATH^. CPMG NMR spectra for 230D7 (red), 230D7 in the presence of 5 μM SPOP^MATH^ (green). The STD spectrum for 230D7 is recorded in the presence of 5 μM SPOP^MATH^. **c**, 230D7 disrupts protein binding between SPOP^MATH^ and PTEN, as measured by *in vitro* pull-down assay. **d**, Concentrations (ng/mL) of 230D7 in BALB/c mice plasma after i.p. administration of 10 mg/kg 230D7. **e**, The Pharmacokinetic parameters of 230D7 after an intraperitoneal dose in BALB/c mice. **f**, The body weight of BALB/c mice treated with different dosages of 230D7 daily for 7 days (n=3 mice each group). **g**, The weight of different organs (heart, liver, spleen, lung, and kidney) of BALB/c mice treated with different dosages of 230D7 daily for 7 days (n=3 mice each group). **h**, Representative histological morphology of H&E-stained tissue sections of BALB/c mice in 230D7-treated or vehicle control groups. **i**, The body weight of NSG mice were measured during the entire pharmacodynamics study of 230D7.

**Extended Data Fig. 5.**
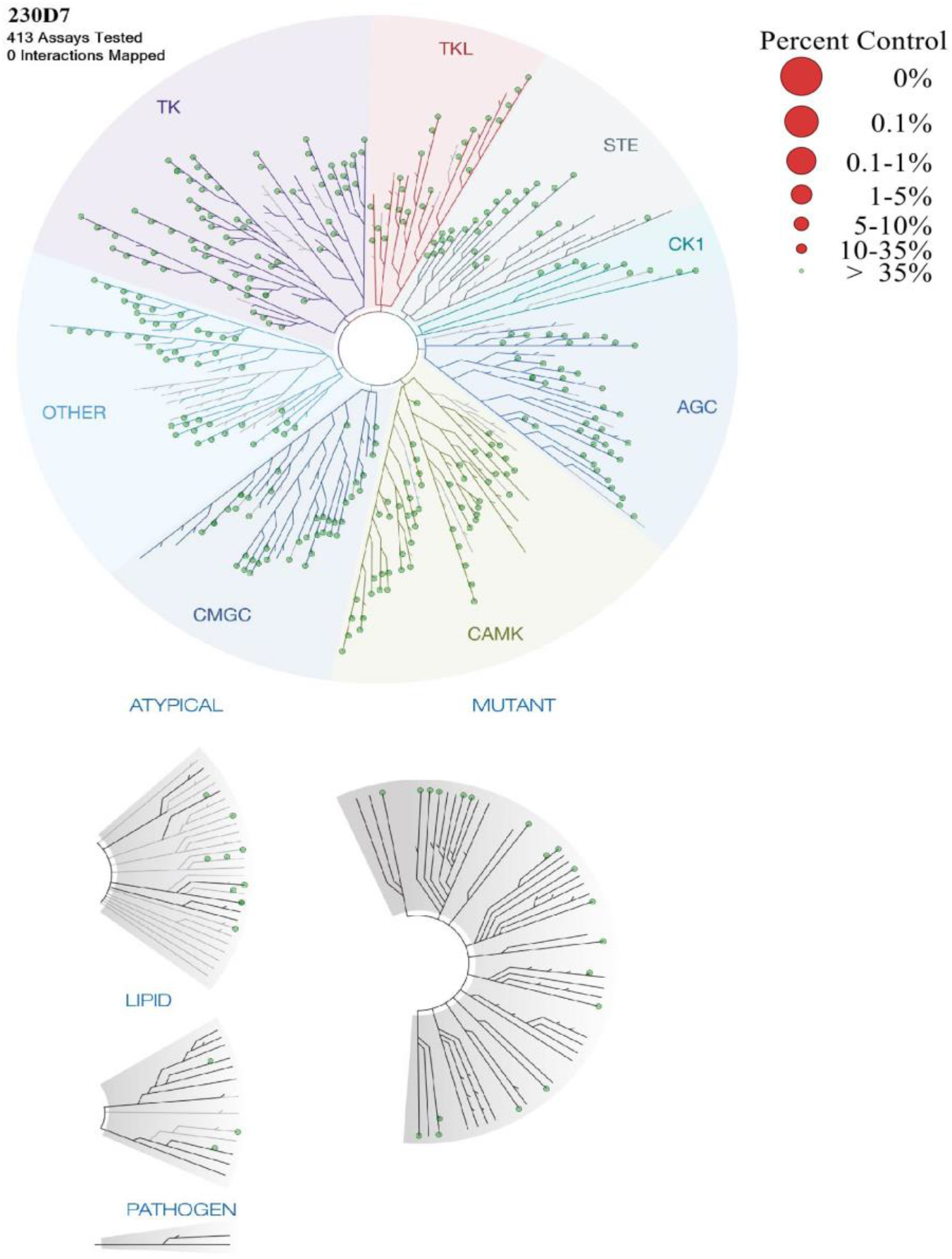
Kinome profiling of 230D7. 230D7 (10 μM) was submitted for a KinaseProfiler (eurofins) to quantify interactions with 413 human wild-type/mutant kinases. The results are displayed as a TREESPOT interaction map. Image generated using TREEspot™ Software Tool and reprinted with permission from KINOMEscan®, a division of DiscoveRx Corporation, © DISCOVERX CORPORATION 2010.

**Extended data Fig. 6.**
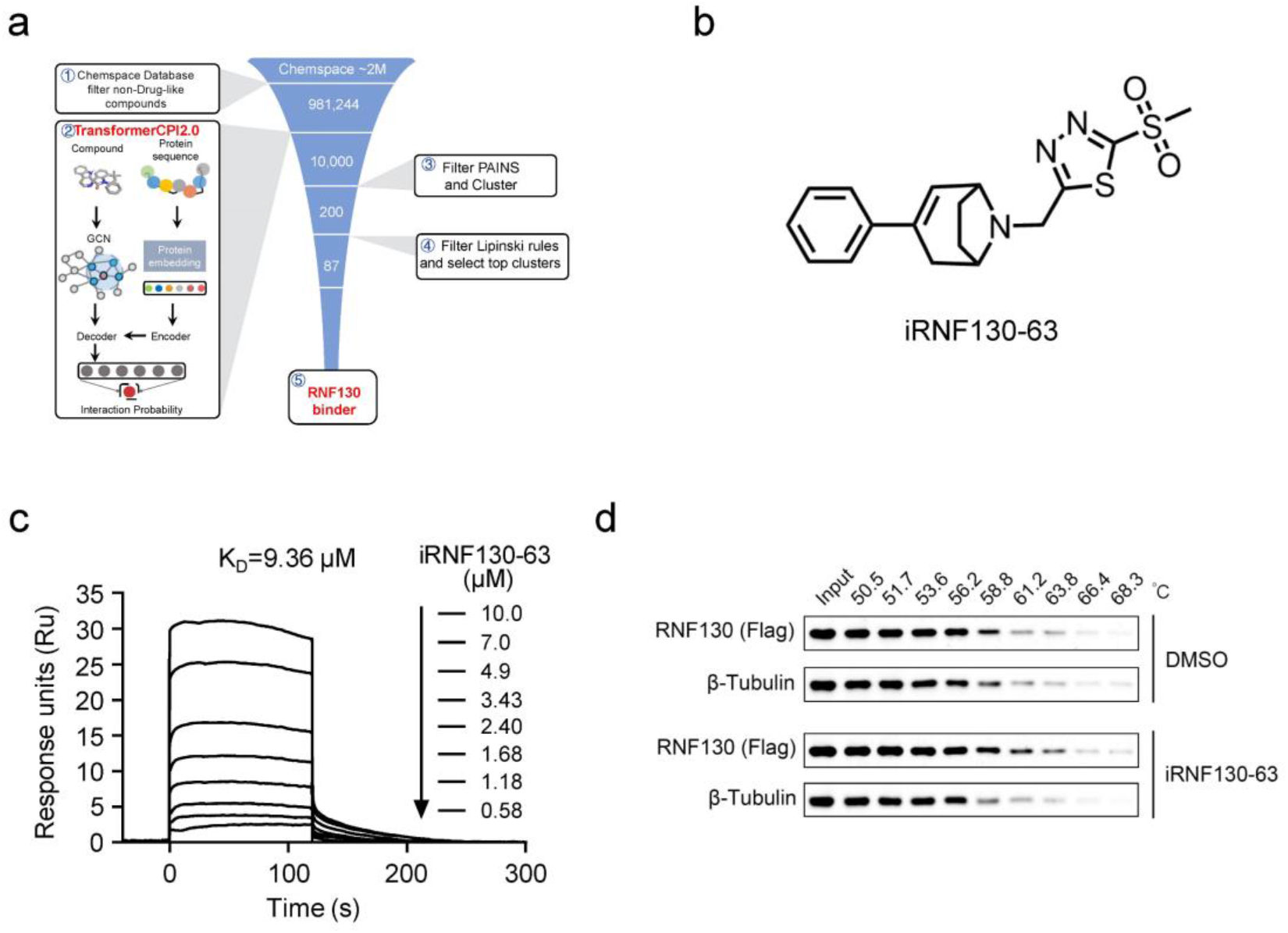
Discovering the chemical binder of RNF130. **a**, The virtual screening procedure of RNF130. **b**, Chemical structure of iRNF130-63. **c**, Surface plasmon resonance analysis examining the direct binding affinities of iRNF130-63 to RNF130. Graphs of equilibrium unit responses versus iRNF130-63 concentrations are plotted. The estimated KD is 9.36 μM. **d**, Representative western blots for the effect of 50 μM iRNF130-63 on the thermal stabilization of RNF130 protein. Cellular thermal shift assay (CETSA) was assayed in 293T cell lysate.

**Extended Data Fig. 7.**
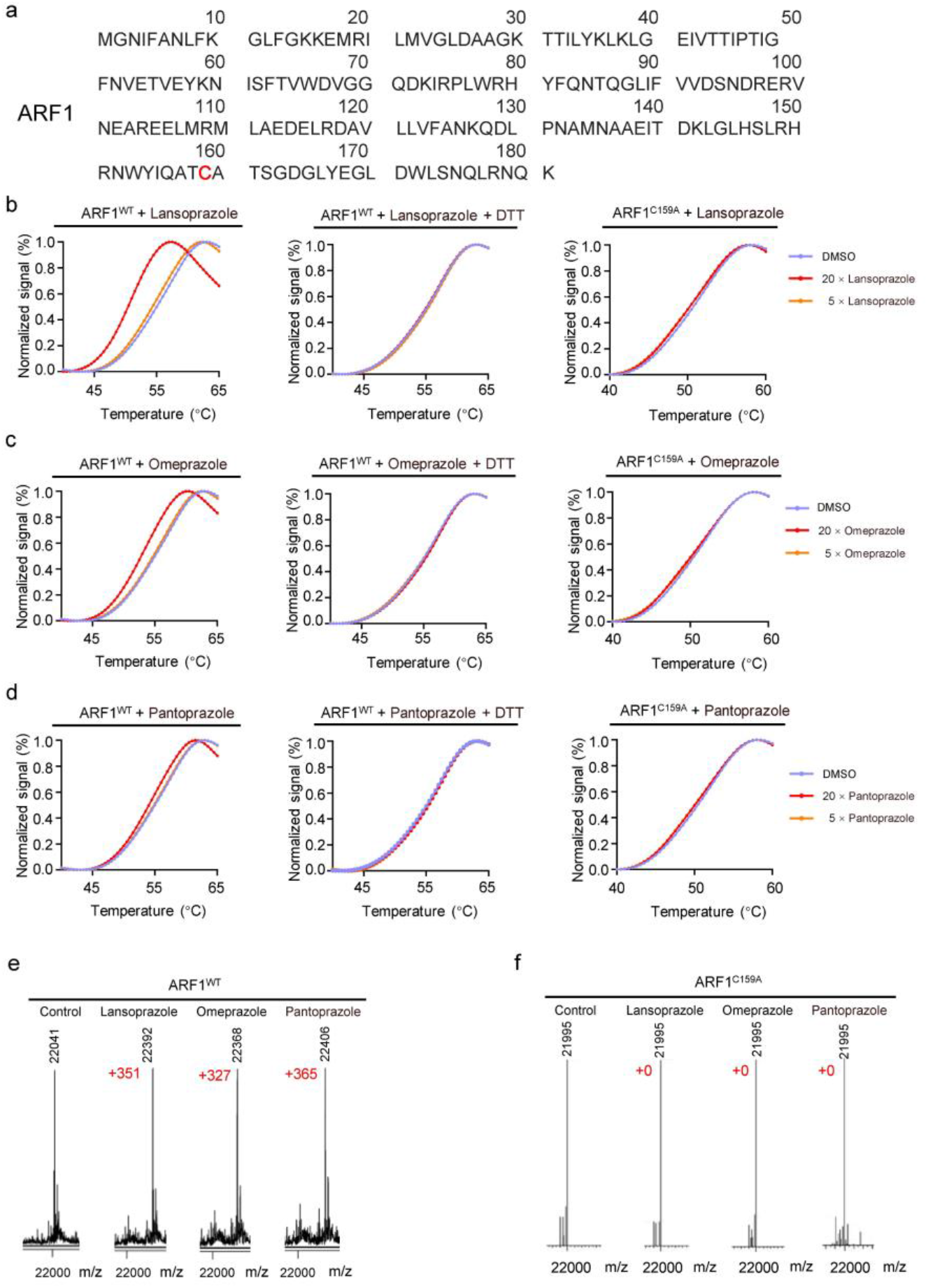
Supplementary data for Fig 5. **a,** Amino acid sequence of ARF1, and C159 (marked red) is the only cysteine residue. **b**, PTS assay of lansoprazole+ ARF1^WT^ (left panel); PTS assay of lansoprazole+ ARF1^WT^+ DTT (middle panel); PTS assay of ARF1^C159A^ + lansoprazole (right panel). **c**, PTS assay of omeprazole+ ARF1^WT^ (left panel); PTS assay of omeprazole+ ARF1^WT^+ DTT (middle panel); PTS assay of ARF1^C159A^ + omeprazole (right panel). **d**, PTS assay of pantoprazole+ ARF1^WT^ (left panel); PTS assay of pantoprazole+ ARF1^WT^+ DTT (middle panel); PTS assay of ARF1^C159A^ + pantoprazole (right panel). **e**, Deconvoluted electrospray ionization mass spectra of ARF1^WT^ in the presence of lansoprazole or omeprazole or pantoprazole. **f**, Deconvoluted electrospray ionization mass spectra of ARF1^C159A^ in the presence of lansoprazole or omeprazole or pantoprazole.

**Extended Data Fig. 8.**
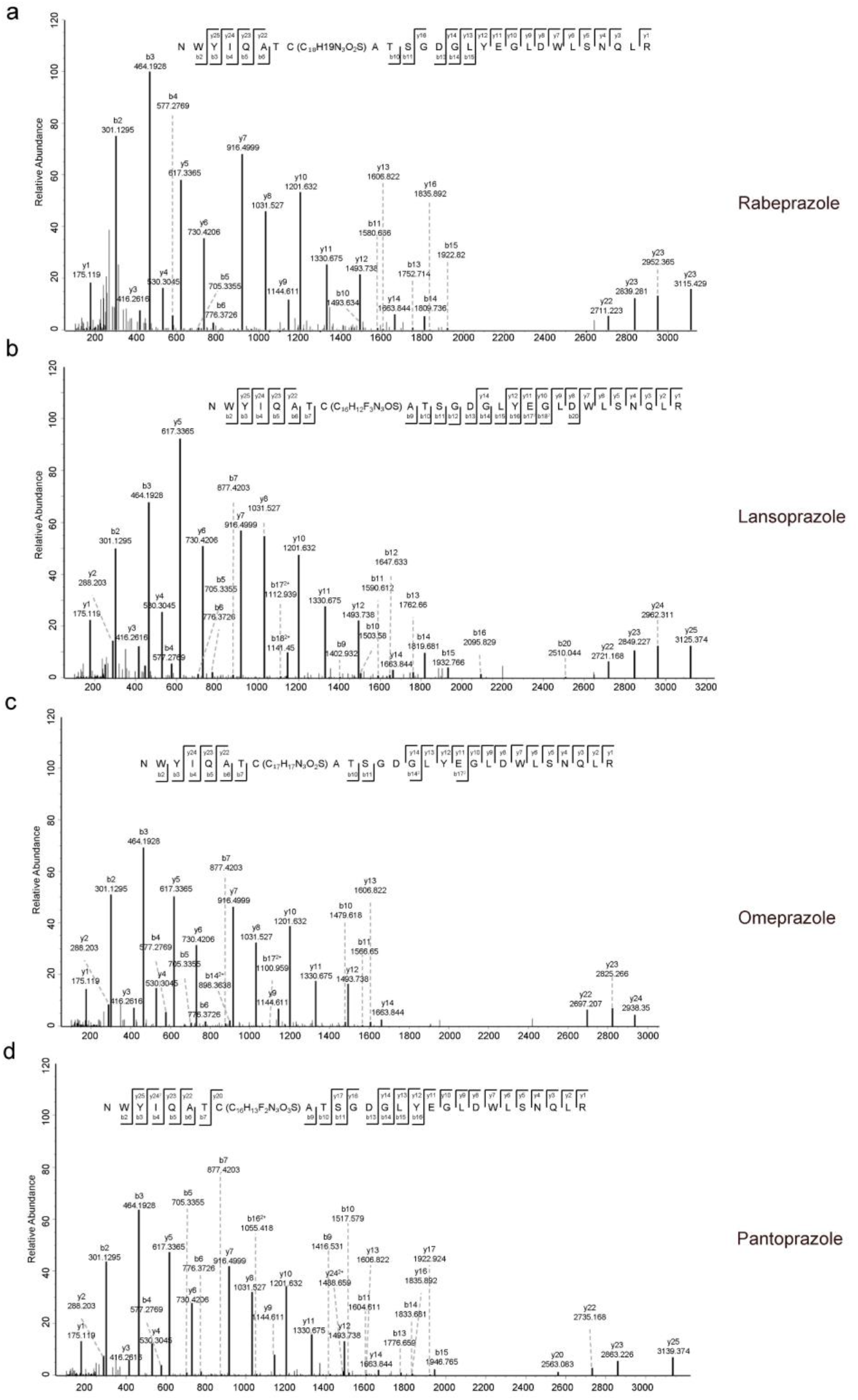
Two-dimensional mass spectra of rabeprazole, lansoprazole, omeprazole, and pantoprazole. **a∼d,** Q-Exactive tandem mass spectra results showed the modified peptide of ARF1, demonstrating that ARF1 was covalently modified by, rabeprazole (a), lansoprazole (b), omeprazole (c) and pantoprazole (d) at cysteine 159.

**Extended Data Fig. 9.**
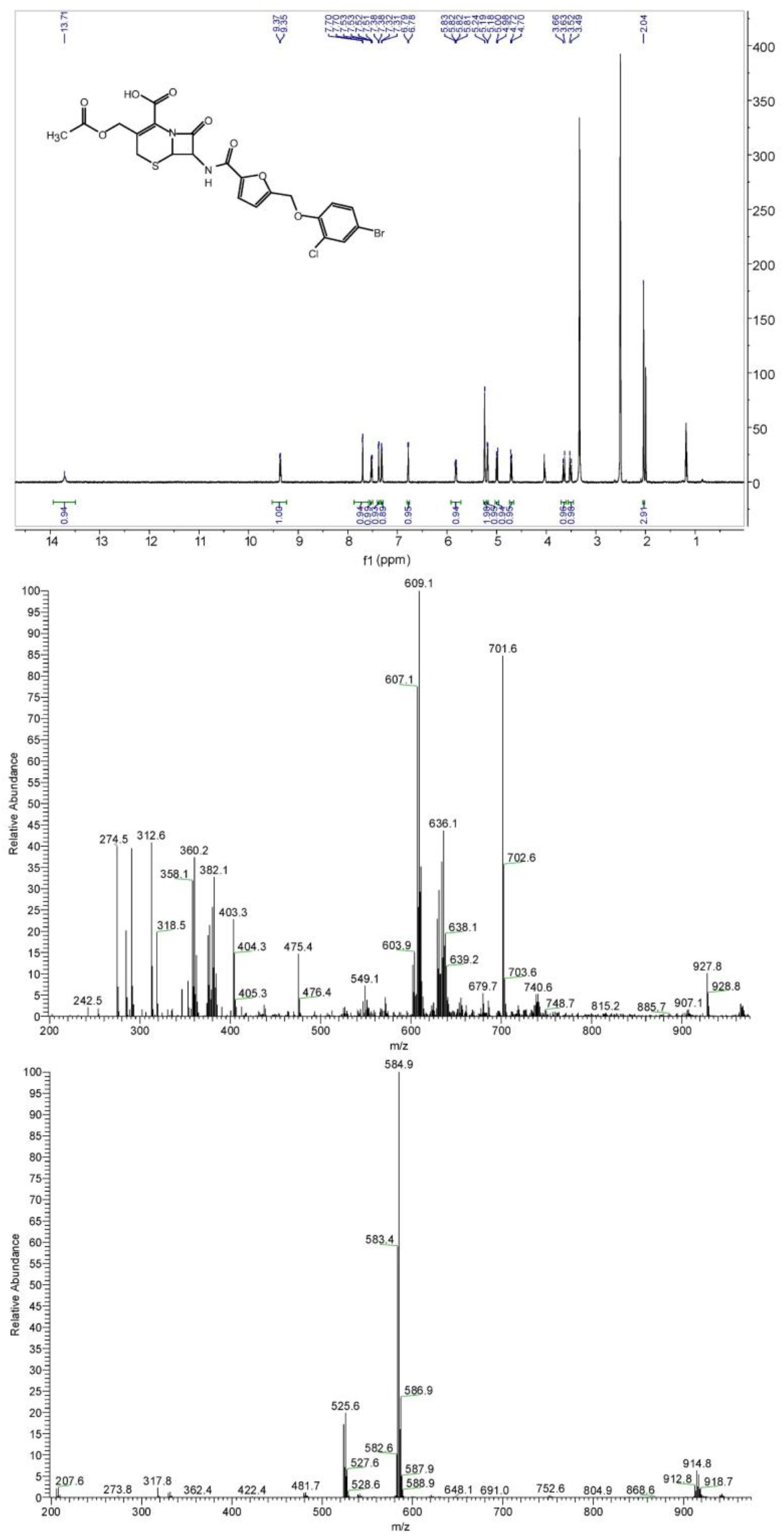
^1^H NMR and ESI-LRMS spectra of 230D7.

## Notes

### Competing Interest Statement

The authors have declared no competing interest.

### Summary of Updates

Release the chemical strutures of SPOP and RNF130 inhibitors.Add a new section titled "Repositioning proton pump inhibitors as anticancer drugs by targeting ARF1".

